# Structural elucidation of recombinant *Trichomonas vaginalis* 20S proteasome bound to covalent inhibitors

**DOI:** 10.1101/2023.08.17.553660

**Authors:** Jan Silhan, Pavla Fajtova, Jitka Bartosova, Brianna M. Hurysz, Jehad Almaliti, Yukiko Miyamoto, Lars Eckmann, William H. Gerwick, Anthony J. O’Donoghue, Evzen Boura

## Abstract

Proteasomes are essential for protein homeostasis in mammalian cells^1-4^ and in protozoan parasites such as *Trichomonas vaginalis (Tv)*.^5^ *Tv* and other protozoan 20S proteasomes have been validated as druggable targets.^6-8^ However, in the case of *Tv* 20S proteasome (*Tv*20S), biochemical and structural studies were impeded by low yields and purity of the native proteasome. We successfully made recombinant *Tv*20S by expressing all seven α and seven β subunits together with the Ump-1 chaperone in insect cells. We isolated recombinant proteasome and showed that it was biochemically indistinguishable from the native enzyme. We confirmed that the recombinant *Tv*20S is inhibited by the natural product marizomib (MZB)^9^ and the recently developed peptide inhibitor carmaphycin-17 (CP-17)^8,10^. Specifically, MZB binds to the β1, β2 and β5 subunits, while CP-17 binds the β2 and β5 subunits. Next, we obtained cryo-EM structures of *Tv*20S in complex with these covalent inhibitors at 2.8Å resolution. The structures revealed the overall fold of the *Tv*20S and the binding mode of MZB and CP-17. Our work explains the low specificity of MZB and higher specificity of CP-17 towards *Tv*20S as compared to human proteasome and provides the platform for the development of *Tv*20S inhibitors for treatment of trichomoniasis.

## Introduction

*Trichomonas vaginalis* (*Tv*), a pear-shaped single-celled protozoan organism, is the etiological agent of trichomoniasis, the most widespread non-viral sexually transmitted disease (STD) worldwide^11,12^. This parasite possesses a single flagellum, which enables its motility, as well as several hair-like structures called pili, facilitating its attachment to host cells. *Tv* also has a complex cytoskeleton that gives it the ability to alter its shape and to traverse host tissues. In women, *Tv* infection can cause vaginitis, while in men it can cause urethritis and prostatitis. Notably, this infection heightens the risk of transmission of HIV and other STDs in both sexes. Current treatment relies on 5-nitroimidazoles^13,14^, however, the emergence of resistant strains poses a significant public health threat due to the lack of alternative treatment options^15,16^. Consequently, novel and effective anti-parasitic compounds are urgently needed. Recently, the critical role of the proteasome in the survival of *Tv* was demonstrated, validating it as a potential drug target for treating trichomoniasis^17^.

The proteasome is a large protein complex that plays a pivotal role in the degradation of cellular proteins. It consists of two main components: the 20S core particle and the 19S regulatory particle. The 20S core particle forms a cylindrical structure consisting of four stacked rings, comprising of two inner rings of seven different β subunits and two outer rings comprised of seven different α subunits. Importantly, the β subunits house the active sites responsible for protein degradation^18,19^. Structural studies using yeast and mammalian proteasomes have greatly improved our understanding of the mechanisms of protein degradation by these enzymes, as well as their interactions with inhibitors and other proteins. To perform structural studies, highly pure and homogeneous samples are required. We and others have previously isolated proteasomes from parasites such as *Plasmodium falciparum* that are sufficiently pure for structural studies^20,21^ but efforts to isolate highly pure proteasome from *Tv* were unsuccessful^8,22^. Therefore, a strategy to make recombinant *Tv* proteasome is needed. Archaeal 20S proteasomes, composed of homo-heptameric rings can be produced by co-expressing the two subunits in *E. coli*^23^. However, the tightly regulated biogenesis pathway of eukaryotic 20S proteasome makes expression in bacteria unfeasible.^24^ The assembly of the human 20S proteasome involves the stepwise incorporation of 14 distinct protein subunits, α1–α7 and β1–β7, assisted by five dedicated chaperones. These chaperones consist of Ump-1 and the heterodimers, PAC1-PAC2 and PAC3-PAC4^25,26^. The 7 α subunits form the α-ring which then serves as a scaffold for the ordered incorporation of β subunits. After dimerization of two pre-assembled half-proteasomes, the final maturation step involves self-cleavage of the β subunit pro-peptides. These N-terminal extensions, present in immature β subunits, are believed to shield their proteolytic activity until formation of the entire 20S proteasome and may also contribute as scaffolds for proteasome assembly^18,27^.

Following maturation, the β1, β2, and β5 subunits contain a threonine residue as the most N-terminal amino acid. This threonine residue can initiate a catalytic reaction through the nucleophilic attack of the carbonyl carbon within the peptide backbone of the substrate. This reaction leads to cleavage of the protein or peptide substrates. The β1, β2, and β5 subunits have distinct substrate specificity preferences and are generally considered to have caspase-like, trypsin-like, and chymotrypsin-like activities, respectively^28^. These three subunits enable the proteasome to cleave a wide variety of substrates at distinct sites allowing to cell to efficiently degrade proteins and maintain cellular protein homeostasis^29,30^.

Proteasome inhibitors bind to the proteasome, hindering its proteolytic activity and impeding protein degradation. This results to the accumulation of proteins within cells, ultimately triggering cell death. Proteasome inhibitors have been shown to selectively kill cancer cells, and three inhibitors (bortezomib, carfilzomib, and ixazomib) have been approved for treatment of multiple myeloma^31^. Additionally, proteasome inhibitors have recently been investigated as potential treatments for parasitic diseases, such as malaria^21,32,33^, Chagas disease and leishmaniasis^34-36^, with one molecule, LXE408, developed by Novartis Pharmaceuticals entering Phase II clinical trials in December 2022. These clinical studies in other parasites support our rationale for developing proteasome inhibitors to treat trichomoniasis.

Carmaphycin B is a marine cyanobacterial metabolite that has been isolated from extracts of *Symploca* sp. collected from Curaçao^10^. This compound is a potent inhibitor of 20S proteasome from mammalian, yeast, nematode and protozoa^10,32,37,38^. Carmaphycin B exhibits potent cytotoxic activity in these cells, and we have synthesized more than 100 analogs that exhibit a range of biological activities. Carmaphycin-17 (CP-17) is one analog that has been shown to have reduced cytotoxicity for human cells while demonstrating significant activity against *Tv*. Additionally, it has shown to be potent against metronidazole-resistant strains.^8^ In topical treatment studies on mice infected with *Tritrichomonas foetus*, a related species suitable for studying vaginal trichomonad infections, CP-17 reduced parasite burden without any noticeable adverse effects. These studies not only validated *Tv* proteasome as a therapeutic target for novel trichomonacidal agents but also highlighted CP-17 as a starting point for further medicinal chemistry studies^8^.

Marizomib (MZB) is a natural product isolated from the marine bacteria *Salinispora tropica*^9^. This non-peptidic proteasome inhibitor is currently in a Phase III clinical trials for the treatment of various types of glioblastoma and stands out as the only proteasome inhibitor capable of readily crossing the blood-brain barrier^39-41^. While MZB primarily targets the β5 subunit of the 20S proteasome, it has been reported to also inhibit the β1 and β2 subunits at higher concentrations^42^. This ability to target multiple subunits of the proteasome may contribute to its potent anti-cancer activity. Compared to clinically approved proteasome inhibitors such as bortezomib and carfilzomib, MZB offers advantages including enhanced stability, bioavailability, and the potential for a broader range of anti-cancer activity.

In this study, we successfully employed the baculovirus expression system to generate recombinant *Tv*20S proteasome. Biochemical comparison with the native enzyme confirmed that all three catalytic subunits of the recombinant enzyme were functional, as they hydrolyzed three different subunit-specific fluorogenic substrates. In addition, a broad-spectrum activity-based probe covalently labelled the catalytic threonine from each subunit. Furthermore, we determined the structure of *Tv*20S in complex with MZB and CP-17. MZB was found to bind to six sites within *Tv*20S that corresponded to three subunits in each β-ring, albeit the electron density was insufficient to fully resolve its binding in the β5 subunit. Conversely, CP-17 bound to four out of six catalytic subunits, corresponding to β2 and β5 from each β-ring. These findings not only offer valuable insights into the structure and function of *Tv*20S proteasome but also shed light on the underlying molecular mechanisms of MZB and CP-17 inhibition.

## Materials and Methods

### Cloning, expression and protein purification

cDNAs encoding all 14 subunits of *Tv*20S proteasome and the proteasome assembly chaperone Ump-1 were synthesized as codon-optimized genes for expression in *E. coli* (Azenta) and cloned into separate pACEBac1 vectors by restriction cloning (**Fig. S1**). The plasmids were transposed to DH10EmBacY cells (Geneva Biotech) and the baculoviruses and protein expression was performed according to manufacturer’s instructions.

The insect cells were centrifuged for 15 min at 2000 x g and the pellet was diluted in Buffer A (50 mM Tris pH 7.5, 150 mM NaCl, 1 mM dithiothreitol, 1 mM EDTA) at a volume corresponding to 4-times the volume of the pellet. Cells were lysed by sonication and the lysate was cleared by centrifugation at 30,000 *g* for 20 minutes. The cleared lysate was loaded onto tandem Streptactin XP columns (QIAGEN), equilibrated in Buffer A. Upon extensive washing with Buffer A, protein was eluted by in the same buffer supplemented with 50 mM biotin. Proteasome containing fractions were pooled, concentrated using ultrafiltration (Amicon) and loaded onto a Superose 6 Increase 10/300 gel filtration column (GE Healthcare) equilibrated with 50 mM Tris pH 7.5, 150 mM NaCl, 1 mM EDTA. 0.5 mL fractions were obtained and assayed for protease activity. Enzymatically active fractions were pooled and concentrated to 1 mg/ml. The native *Tv*20S proteasome was purified from frozen pellets of *Tv* parasites as described previously^22^.

### Proteasome activity and inhibition assays

The β5 proteolytic activity of the 20S proteasome was measured in insect cell extracts and in fractions isolated from chromatography columns using the fluorogenic substrate, Suc-LLVY-amc (Cayman). For enzyme characterization studies, substrates specific for Tv20S β1, β2, and β5 subunits, namely Ac-RYFD-amc, Ac-FRSR-amc, and Ac-GWYL-amc, were used as described previously.^22^ These substrates were custom synthesised by GenScript, NJ. For inhibition studies, 1 μM recombinant *Tv*20S was incubated with 50 μM MZB, 50 μM CP-17 inhibitor, or 0.5% DMSO (vehicle control) for 1.5 hours at room temperature. The inhibition of individual Tv20S subunits was confirmed using the subunit specific fluorogenic substrates. Kinetic assays in 384-well plates were performed using 5 nM Tv20S in 50 mM HEPES pH 7.5 with 50 μM of Ac-RYFD-amc, 50 μM of Ac-FRSR-amc, and 50 μM of Ac-GWYL-amc, in a final volume of 30 μL per well. All assays were performed in triplicate wells on 384-well black plates (Nunc) at 37°C in a Synergy HTX Multi-Mode Microplate Reader (BioTek, Winooski, VT) with excitation and emission wavelengths of 360 and 460 nm, respectively.

### Protein Gels and Active Based Probing

Native *Tv*20S (n*Tv*20S), recombinant *Tv*20S (r*Tv*20S) and r*Tv*20S-inhibitor complex were diluted with 50 mM HEPES pH 7.5 then mixed with 2 μM MV151 (R&D Systems #I-190). After MV151 addition, samples were incubated at RT for 16 hours. For denaturing gels, samples were mixed with 4X Bolt LDS sample buffer (Thermo) containing 250 μM dithiothreitol, heated at 100°C for 5 minutes and loaded into a NuPAGE 12% Bis-Tris gel (Thermo). PageRuler Plus pre-stained protein ladder (Thermo) was included on each gel. Gels were run with 1X MOPS SDS buffers (Invitrogen) at 130V. For native gels, samples were mixed with 2X Novex Tris-glycine native sample buffer and loaded into NuPAGE 3-8% Tris-glycine gels (Invitrogen) with NativeMark unstained protein standard (Thermo). Gels were run at 100V with Novex Tris-glycine running buffer (Invitrogen). All gels were imaged on Bio-Rad ChemiDoc XRS+ at 470 nm excitation 530 nm emission for MV151 probe visualization and silver stain or coomassie brilliant blue stain.

### Preparation of cryo-EM grids and data acquisition

The recombinant purified *Tv*20S (1.4 μM) was mixed with 1.25 mM of either MZB or CP-17 to achieve a final concentration of 50 μM inhibitor. The mixture was prepared in a solution of 50 mM HEPES pH 7.5 and incubated at room temperature for 1 hour. After the incubation period, the sample was cooled on ice. Simultaneously, the cryo-EM Quantifoil R2/1 300-mesh copper grids (EM Sciences, Prod. No. Q350CR1) were glow discharged at 15 mA for 30 seconds to enhance their surface properties. 4 μL of proteasomes-inhibitor complex at a concentration of 1 mg/ml was transferred to the grid. The grids were blotted at -5 power for 5 s in FEI Vitrobot Mark IV (Thermo Fisher Scientific) at 4°C and 100% humidity and immediately frozen in liquid ethane. Immersion freezing and screening of cryo-EM data and all data acquisition were performed at the UmeÅ Core Facility for Electron Microscopy, Sweden. Screening was performed using a 200 kV Glacios system (Thermo Fisher Scientific) equipped with a Falcon 4i direct electron detector. 7 983 images of *Tv*20S with MZB and 10 037 images of *Tv*20S with CP-17 were acquired with pixel size of 0.7 Å with an exposure of 40 electrons per Å^2^ on Titan Krios (Thermo Fisher Scientific) at 300 kV using a Falcon 4i direct electron detector.

### Cryo-EM imaging

The data for *Tv*20S complexed with MZB was processed using a cryoSPARC (v4.0.1). Images were imported and corrected using default settings for patch motion and patch contrast transfer function (CTF). Particles were selected using a blob picker (with a particle search size of 190 Å), extracted with a box size of 440 pixels, and classified using a 2D classification job. For *Tv*20S-MZB initial 100 2D classes were sorted using 2D classification. Particles and exposures were reduced to 6135 particles using a manual curation job. Subsequent 2D classification revealed that of the 916 422 good particles, only 2% of the 2D classes contained the entire proteasome. After several rounds of 2D classification to remove unwanted particles, the subset of 13 933 particles (side views only) was used for an *ab initio* reconstruction and further homogeneous refinement. The resolution was improved by 3D classification and removal of poor 3D classes and further reiterations including 2D classifications. The C2 symmetry was applied in 3D homologous refinement steps for generation of final model. The cryo-EM map (GSFSC) achieved a final resolution of 2.86 Å, utilizing 14 257 particles.

The cryo-EM 3D map for *Tv*20S complexed with CP-17 was generated in a similar manner. Images were imported in small batches and then pooled, motion-corrected, and CTF-corrected in four batches, each containing approximately 2000 images. Particle picking was performed using well-defined *Tv*20S-MZB particles as templates. 2D classifications were also conducted in four batches, and subsequently, visually well-defined particles were selected (528 679), reclassified, and filtered using both 2D and 3D classification methods. After a several rounds of further selection processes, just 80 145 particles were retained for the final 3D map generation. The resulting cryo-EM map of *Tv*20S complexed with CP-17 exhibited a final resolution estimate of 2.60 Å, as determined by GSFSC analysis. For overview of cryo-EM data processing workflow please refer to **Fig. S2**.

### Model building and refinement

A high-resolution proteasome structure of *Leishmania tarentolae* (PDB ID: 7ZYJ) was used as an initial template to build the model.^43^ The initial model was built by alignment of *Tv*20S the structures predicted by AlphaFold onto the 7ZYJ template. The *Tv*20S-MZB model was fitted to density using ChimeraX^44^ and refined by rigid body refinement and real space refinement in Coot 0.9.6 EL^45^ and structure optimisation using ISOLDE package^46^ in ChimeraX. Representative images of the structures and maps were generated using ChimeraX and Pymol. *Tv*20S-CP-17 structure which was refined in the similar manner as the *Tv*20S-MZB structure using the *Tv*20S-MZB as a starting model. The *Tv*20S-MZB structure was deposited to world protein data bank with PDB ID: 8OIX and similarly the *Tv*20S-CP-17 structure was deposited as PDB ID: 8P0T (**SI Table 1**).

## Results

### Preparation of the Tv20S

The genome of *T. vaginalis* G3 is available from trichdb.org and all 14 subunits of the *Tv*20S proteasome were identified by alignment studies with the human constitutive proteasome (**Fig. S3**). From these studies, we identified the putative β7 subunit as A2F3X4 (Uniprot ID) which in the human and yeast proteasome has been shown to be the last subunit incorporated into the α-ring/β-ring half-proteasome^47,48^(**Fig. 1**). A sequence encoding a C-terminal twin-strep tag was added to the β7 subunit as this was expected to have minimal effect of the assembly of the proteasome complex. In addition, the *Tv* genome was searched for chaperone proteins with homology to PAC1–PAC2, PAC3–PAC4 or Ump-1 that are known to play a role in proteasome assembly in human cells^24,49^. A homolog of human Ump-1 (Uniprot A2FJW0) with 21% sequence identity was identified (**Fig. S3**). No homologs of PAC1–PAC2, PAC3–PAC4 were found.

**Fig. 1:**
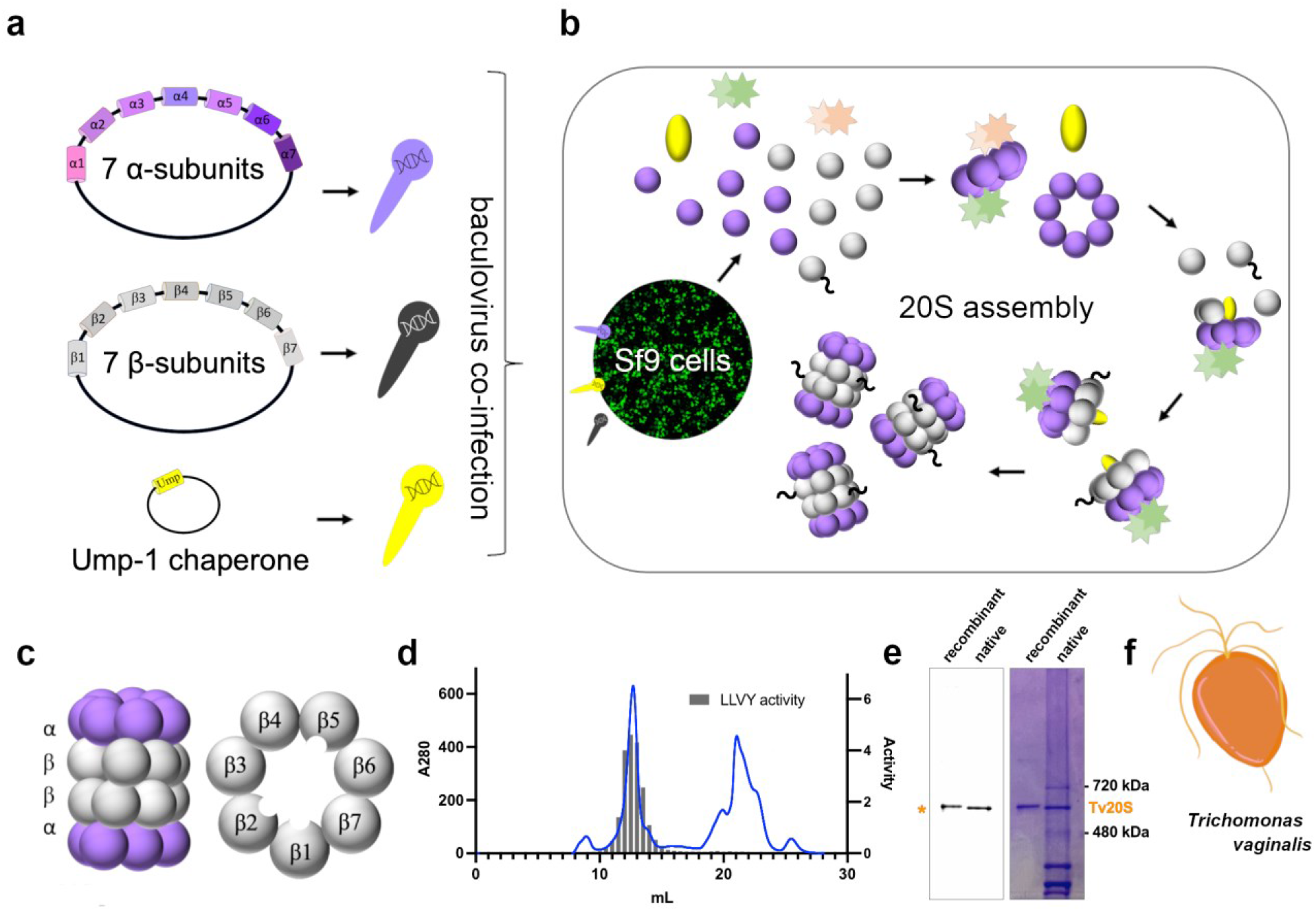
Cloning, Recombinant Expression, and Purification of Tv20S Proteasome. **(a)** Cloning of three vectors containing either 7 α-subunits, 7 β-subunits or Ump-1 into baculovirus. **(b)** Co-infection of SF9 cells with baculovirus for proteasome expression and assembly. **c**) Side view structure of the 20S proteasome complex showing two β rings sandwiched between two α rings. Planer view of the β ring showing the three catalytic subunits located within the central tunnel of the proteasome. **d**) Final purification step using Superose 6 chromatography. Absorbance at 280 nm (blue line) and proteasome activity assessed using a proteasome-specific fluorogenic substrate Suc-LLVY-amc (grey bar chart). MV151 and Coomassie blue-stained native PAGE showing the purity of the recombinantly purified proteasome complex compared to native Tv20S isolated from *T. vaginalis*. F) Cartoon representation of *T. vaginalis* at 100x magnification.

Three baculoviruses were prepared, one bearing seven α genes, one with seven β genes, and a third with the Ump-1 chaperone gene (**Fig. 1a**). Insect cells were simultaneously infected with the three different baculoviruses (**Fig. 1b**) and the resulting recombinant protein in the cell lysate was enriched on a Strep-tag XT column and then further purified on a Superose 6 column. Only fractions containing catalytically active protein were selected following assays with the Suc-LLVY-amc substrate (**Fig. 1d**). A protein yield of ∼1 mg per liter of cell culture was achieved.

To compare r*Tv*20S with the native *Tv*20S (n*Tv*20S) proteasome, both were incubated with a fluorescently labeled inhibitor probe, MV151. Gel electrophoresis showed that MV151 bound to the catalytic subunits of Tv20S, resulting in a band with a molecular weight of approximately 690 kDa for both the native and recombinant proteasomes. The recombinant enzyme appeared slightly larger on the gel due to the presence of a 28 amino acid twin Strep-tag on each β7 subunit, adding approximately 6 kDa in size (**Fig. 1e**). Coomassie Blue staining confirmed the high purity of the recombinant preparation compared to the native proteasome that was enriched from a *Tv* cell extract using sequential steps of ammonium sulfate precipitation, size exclusion chromatography and anion exchange chromatography. These findings demonstrate that the recombinant *Tv*20S is highly pure, enzymatically active and forms a complex of similar size to the native enzyme, making it suitable for biochemical and structural studies.

### Characterization of Tv20S-inhibitor complex

We evaluated the function of the individual catalytic subunits of recombinant proteasome using a combination of inhibitors, probes, and fluorogenic substrates. Recombinant *Tv*20S was labeled with the fluorescent probe MV151 which binds irreversible to the catalytic Thr of β1, β2 and β5. Each protein can be visualized on a denaturing gel (**Fig. 2a**). This revealed that β2 and β5 subunits were labeled more efficiently than the β1 subunit, as we have seen previously with the n*Tv*20S.^22^ When 1 μM r*Tv*20S was preincubated with 50 μM of CP-17 (**Fig. 2b**) and then incubated with MV151, the β2 and β5 subunits could not be labeled while the β1 subunit labelling was the same as the uninhibited enzyme (**Fig. 2c**). This revealed that CP-17 binds to β2 and β5 which then prevents labelling of these subunits by MV151. In parallel, we pre-incubated r*Tv*20S with 50 μM MZB and then incubated with MV151, but no bands were visible. This showed that MZB binds to all three catalytic subunits of the proteasome (**Fig. 2c**).

**Fig. 2:**
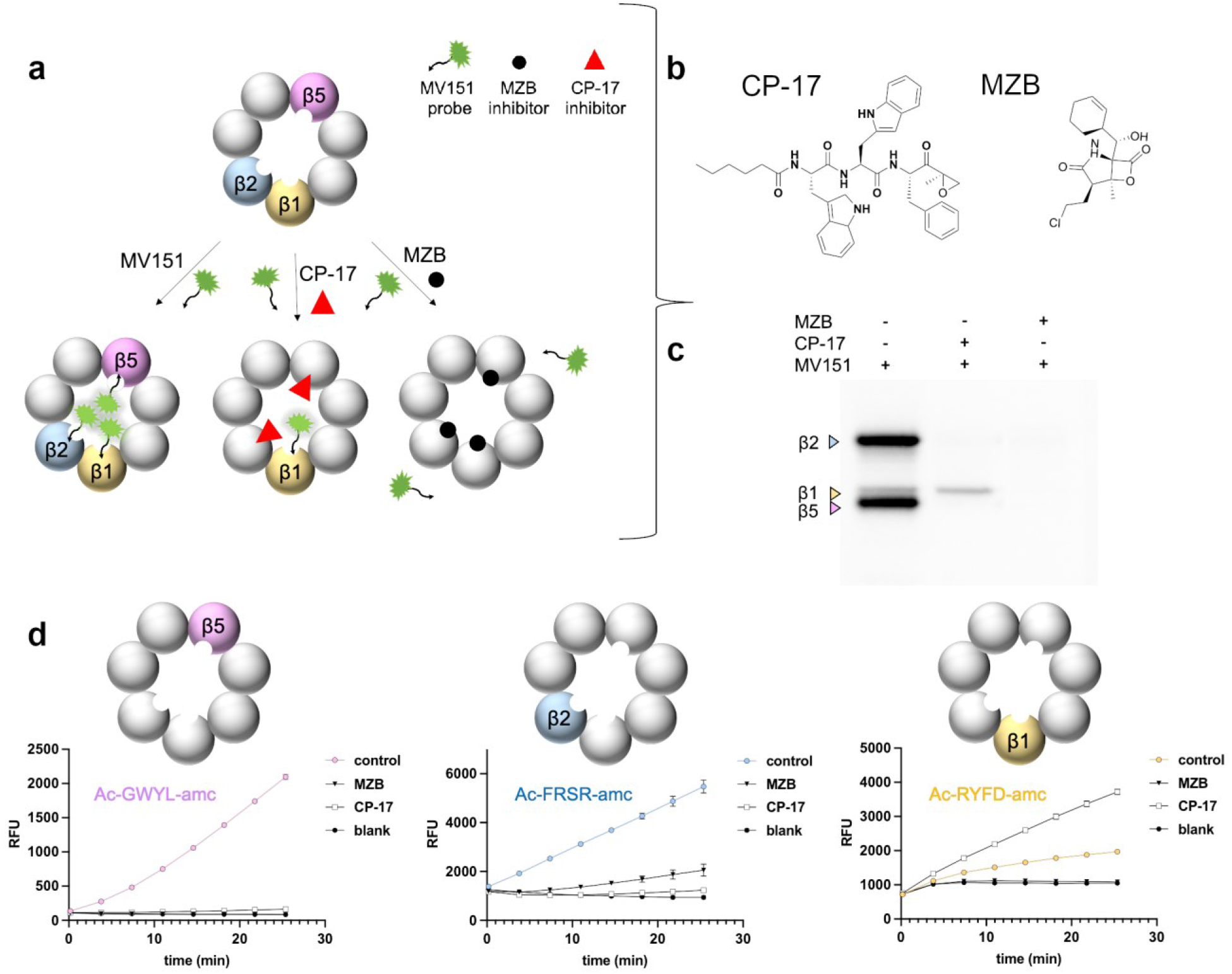
Characterization of the proteasome-inhibitor interaction and inhibition kinetics. **a**) The figure illustrates the schematic representation of active sites during inhibition by a MZB or CP-17, followed by visualization through fluorescent MV151 probing. The inhibitor is depicted binding to specific active sites, thereby blocking enzymatic activity. Subsequent visualization techniques provide insights into the binding interaction and potential conformational changes induced by the inhibitor. **b**) Chemical structure of MZB and CP-17. **c**) rTv20S was first incubated with DMSO, 50 µM CP-17 or 50 µM inhibitor for 1 hour, and then incubated with 2 µM MV151 for 16 hours and the catalytic subunits were imaged on a denaturing gel at 470 nm excitation 530 nm emission. **d**) Screening and validation of inhibitor specificity using subunit-specific proteasome substrates.

Functionality of r*Tv*20S was further confirmed through enzyme kinetic assays using subunit-specific fluorescent substrates. These substrates were rationally designed such that each are cleaved by only one subunit of *Tv*20S.^22^ The substrate for β5, consisting of Ac-GWYL-amc, was efficiently cleaved by r*Tv*20S, however, after pre-incubating with 50 μM CP-17 or MZB, the activity was inhibited by >97% (**Fig. 2d**). Recombinant *Tv*20S was also assayed with the β2-specific substrate, Ac-FRSR-amc and activity was reduced by 76% and 96% in the presence of MZB and CP-17, respectively (**Fig 2e**). β1 activity was detected using Ac-RYFD-amc but pre-incubation with CP-17 led to a 3-fold increase in activity, while MZB completely inhibited its activity. Activation of β1 in the presence of CP-17 was previously observed for n*Tv*20S.^22^ In summary, these probe-labelling and enzyme activity assays reveal that r*Tv*20S is enzymatically identical to n*Tv*20S and therefore, validates the use of the recombinant enzyme for all future biochemical studies.

### Structural characterization of the proteasome from Tv

To obtain atomic models of structures with inhibitors bound to *Tv*20S, we mixed r*Tv*20S proteasome with an excess of CP-17 or MZB and subjected it to cryo-EM. Initial data analysis revealed that the recombinant sample with MZB contained a significant amount of unassembled proteasome (**Fig. S2a**). During the 2D classification of the collected dataset, only 2% of the particles corresponded to the 20S proteasome, while the remaining ring particles mostly consisted of half-proteasome complexes that are made up of a single α and β ring.

Despite this, we were able to obtain a 2.86 Å cryo-EM map of the 20S proteasome, which was used to build the initial atomic model of the r*Tv*20S with MZB (PDB ID: 8OIX). For r*Tv*20S bound to CP-17, we performed a second round of gel filtration. Each fraction obtained was then analyzed for the presence of fully assembled *Tv*20S using negative-stain electron microscopy. This procedure yielded approximately 80% of all particles to be assigned as to the fully assembled *Tv*20S proteasomes. The data collected from this sample led to the cryo-EM reconstruction and determination of the atomic structure of *Tv*20S with CP-17 (PDB ID: 8P0T).

The initial *Tv*20S atomic model was built by leveraging the AlphaFold models, which were superimposed with the existing 20S structure of *Leishmania tarentolae* 20S proteasome (PDB ID: 7ZYJ) and fitted to the cryo-EM maps. Subsequently, the model was refined and rebuilt based on these maps. Unsurprisingly, *Tv*20S exhibits a shape and fold consistent with the characteristic structure of 20S proteasomes. *Tv*20S displays C2 symmetry, with two sets of 14 subunits arranged in the conventional α1–α7, β1–β7 / β1–β7, α1–α7 rings configuration.

Despite the homology among the individual chains, there are variations in their arrangement. The packing of the *Tv*20S proteasome differs from other proteasomes with known structures (e.g., RMSD = 2.729 Å for Cα atoms in structural alignment with the *L. tarentolae* 20S proteasome). It is worth noting that, in these structures, the *Tv*20S sequence (3,167 residues) was mapped onto the cryo-EM maps, achieving 96.8% coverage for *Tv*20S/MZB and 96.6% for *Tv*20S/CP-17 structures. This includes well-defined regions, such as the active site pockets, which allowed for the modeling of inhibitor molecules in both structures.

### Catalytically active sites β1, β2 and β5, and inhibitors MZB and CP-17

All three active sites in *Tv*20S are defined by a conserved catalytic triad Thr1, Asp17 and Lys33. In addition, other well-conserved residues, such as Asp168, Ser131 and Ser170 in β1, are required for efficient catalysis and active site structural integrity ^50^. In the Tv20S-MZB dataset, the electron map within the β1 active site exhibited clear density extending beyond Thr1, providing sufficient coverage for the inhibitor MZB. As a result, the atomic model of MZB was successfully built and fit accurately into the cryo-EM map (**Fig. S4**). Conversely, no density was observed within the β1 active site for CP-17 in the other dataset. This is consistent with the biochemical data (**Fig. 2d**), which demonstrated that MZB inhibits β1 activity while CP-17 does not.

The second active site, β2, can accommodate both MZB and CP-17 inhibitors, as clearly shown by the cryo-EM maps in those regions (**Fig. S4 and S5**). MZB adopts a conformation like that observed in the β1 site. In both cases, the cyclohexenyl moiety of MZB and the phenyl residue of CP-17 occupy a deep pocket that is shared by residues, including Lys33 and a loop composed of Ala46, Ala49, Glu31, and Asn52 (**Fig. 3f**). This pocket is commonly referred to as the S1 pocket of the enzyme because it serves as the binding site for the P1 amino acid of the substrate, which is on the N-terminal side of the scissile bond. For CP-17, the α,β-epoxyketone group is covalently bound to Thr-1 forming a morpholino derivative. The adjacent phenyl side chain of phenylalanine interacts with X, Y and Z. Beyond the S1 pocket, the P2 tryptophan of CP-17 interacts with the enzyme through a proximal indole ring that resides on the hydrophilic ridge of the shallow S2 pocket, while the indole ring of the P3 tryptophan is buried and fits well within the deep S3 pocket. The S3 site is characterized by the presence of aliphatic chains, including Val28 and six alanine residues (Ala20, Ala22, Ala27, Ala124, Ala130, Ala132). Additionally, Glu31 and Asp126 line the S3 pocket, creating an acidic environment. The N-terminal capping group of CP-17 is a pentyl chain, which is orientated in a shallow S4 pocket.

**Fig. 3:**
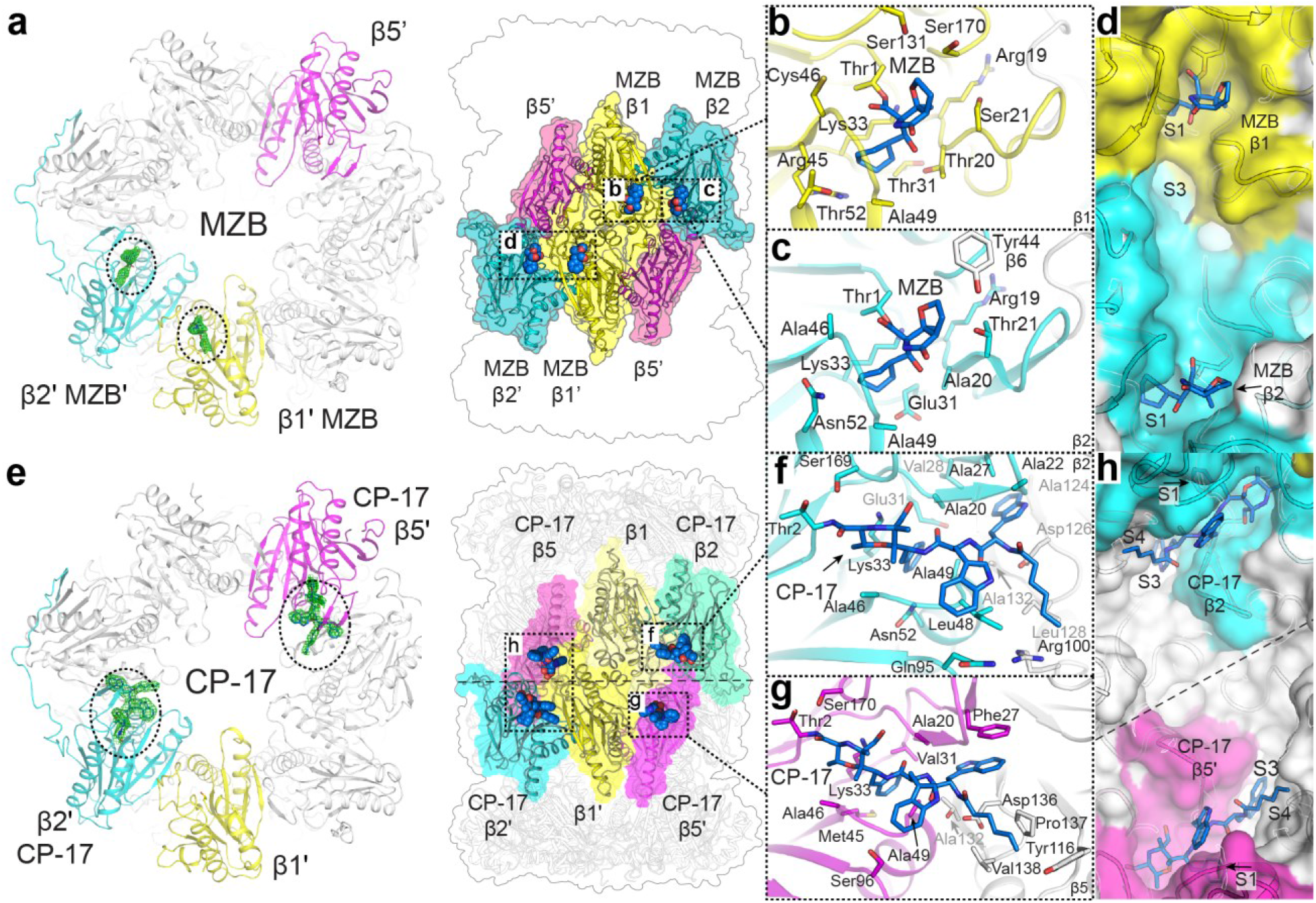
Cryo-EM structures inhibitors MZB and CP-17 bound active sites of *Tv*20S proteasome. The atomic models of *Tv*20S with covalent inhibitors **a** MZB or **e** CP-17. The panels **a** and **e**, the cryo-electron map highlights the inhibitors in active sites (shown as green mesh), while catalytic units are depicted in different colors (β1 yellow, β2 cyan and β5 magenta). The electron map for MZB was observed only in two catalytic sites β1 and β2 (panels **b&c**), on the other hand the electron map for CP-17 was observed only in catalytic sites β2 and β5 (panels **f&g**). These four panels (**b**,**c**,**f**,**g**) show a stick representation of inhibitors (orange and blue) covalently linked to active sites threonine residues. Amino acid residues within a reach of non-covalent interaction (up to 4 Å) are shown as sticks. Panels **d** and **h** show depict the surface representation and structural orchestration of the active site pockets and their connection within *Tv*20S proteasome with the inhibitors bound. The dashed line outlines the C2 symmetry.

In the catalytic site of β5, some density was observed in the cryo-EM map that corresponded to MZB, however unlike the observations made for the β1 and β2 sites, MZB in β5 was not sufficiently defined to allow for a satisfactory fit. Even though MZB is known to bind and inactivate the β5 catalytic activity, it could not be included in the final model. In contrast, the density of CP-17 in the β5 site was clearly defined. The ligand was positioned in a highly similar conformation to that observed in the β2 site. The shapes of the β5 and β2 sites are very similar, although there are differences in the amino acid composition of the S1-S4 pockets (**Fig. 3**). The α,β-epoxyketone group was covalently bound to Thr-1, the P1 phenylalanine is located in the S1 pocket, which is lined with (Lys33, Ala46, and Ala69). In contrast to the β1 and β2 sites, the S1 of β5 harbors Val31 and Met35. Moving forward, the P2 indole ring is positioned on the ridge of S2, formed by the loop between Ala46 and Ala49, and is enclosed by Ser96. The P3 indole is anchored in the relatively deep S3 pocket, defined by phi-phi interaction with the residue Phe27. Finally, the pentyl group of CP-17 sits in a well-defined shallow pocket formed by the hydrophobic parts of four residues belonging to the neighboring β6 chain (Tyr116, Asp136, Pro137, and Val138).

## Comparison of human and trichomonas active sites to reveals potential hotspots for inhibitor design improvements

At first glance, the overall fold of human constitutive proteasomes and *Tv*20S appears identical. However, upon structural alignment, the overall root mean square deviation (RMSD) is calculated to be 2.638 Å, which is higher than anticipated given the conservation of individual subunits (**Fig. 4a**). This disparity can be attributed to the slightly different packing of *Tv*20S, leading to a less optimal overall alignment. When aligning the individual subunits, the RMSD values decrease accordingly as the aligned sequences exhibit a range of 25-51% identity, with conserved regions in the active sites (Table S2). Despite this relatively high sequence identity, the fine structural variances within the active pockets present opportunities for designing selective inhibitors targeting *Tv*20S, as seen in the case of CP-17. CP-17 inhibits the Tv20S β5 subunit with a 10-fold higher potency than the equivalent subunit of the human constitutive proteasome^8,51^. In the human 20S proteasome, several amino acids pose steric hindrances, resulting in clashes when hypothetically positioning the CP-17 inhibitor within the active sites based on the alignment of Tv20S and the human proteasome.

**Fig. 4:**
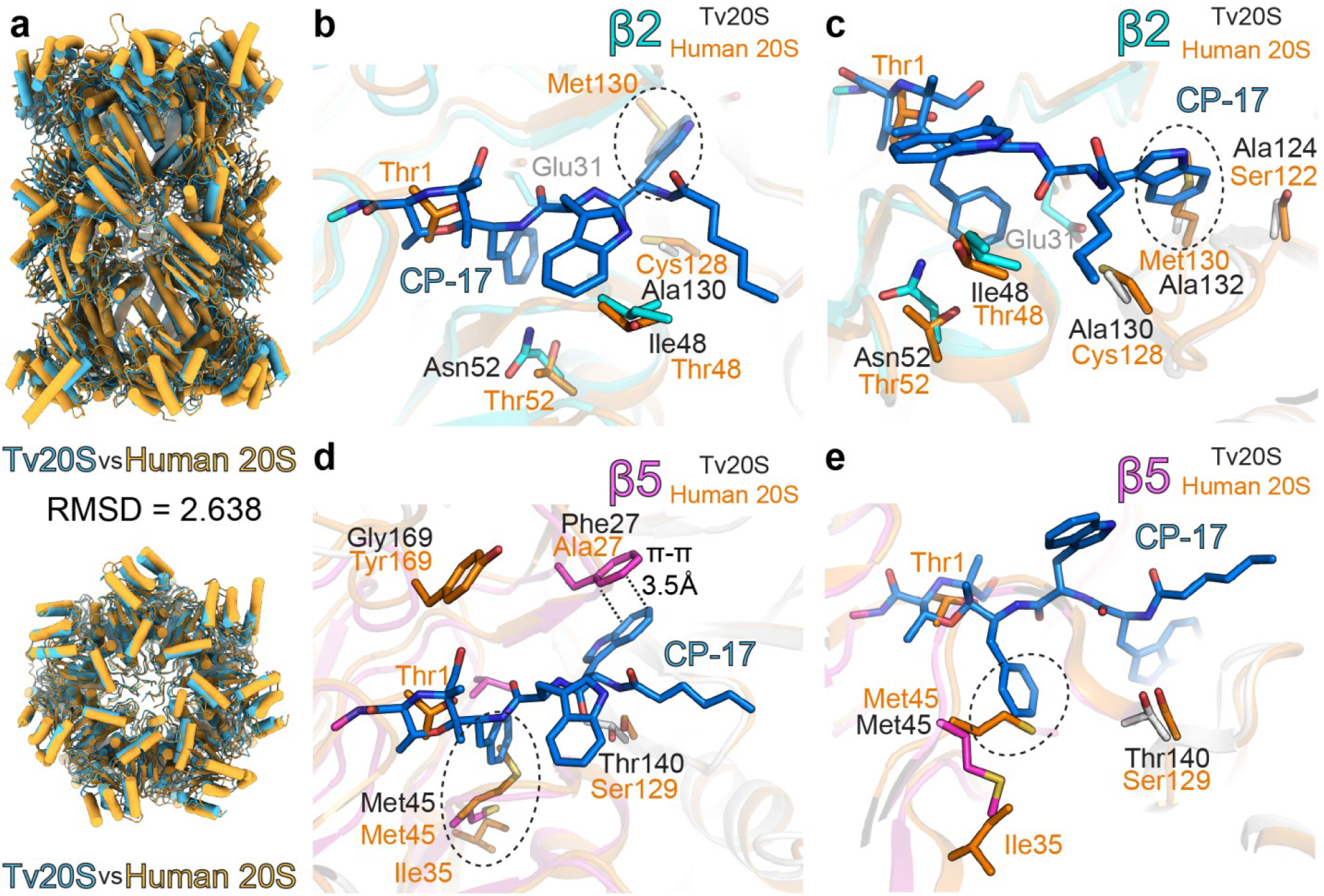
Comparison of *Tv*20S and human 20S structures: The panel **a** comparison of human (orange) and *Trichomonas vaginalis* (light blue) 20S proteasomes. The further panels show structural overlay of human 20S and covalent inhibitor CP-17 (blue sticks) in the active site pockets of β2 (panels **b&c**) and β5 (panels **d&e**) of *Tv*20S. The residues responsible for major structural differences in these active sites are shown as sticks (human-orange, *Tv*20S β2 cyan and β5 magenta, other subunits shown in white). The predicted clashes between CP-17 and amino acids from human 20S are encircled with dotted line. In panels **b&c**, Met130 of human β2 is in clash with one of the indole rings of CP-17, whilst in *Tv*20S this moiety is accommodated in a deep hydrophobic pocket formed by Ala132 and Ala130. Similarly, in panels **d&e**, the pocked formed by Met45 of *Tv*20S β5 allows sufficient space for phenyl ring of CP-17. In this location human β5 Met45 is pushed by Ile35 towards the active site, forming a potential barrier for CP-17.

In the β2 active site much of the residues are conserved but in the S3 pocket where the second indole ring is positioned human 20S contains bulkier residues, such as Met130 and Ser122, in contrast to the significantly smaller alanine residues found at these positions in the *Tv*20S β2 site (**Fig. 4b-c**). The S3 pocket of β5 subunit of *Tv*20S contains Phe27, which forms a favorable π-π interaction with the indole ring and represents the sole significantly different residue in this site. Conversely, the S1 pocket of the human active site contains Met45 and Ile35, which may obstruct the binding of the phenyl ring (**Fig. 4e-f**). It is plausible that Met45 from *Tv*20S exhibits greater mobility, moving away from the approaching ligand and not being hindered by Ile35, while the π-π interaction with Phe27 enhances the binding of the ligand in the case of *Tv*20S proteasome. This differential binding mechanism allows CP-17 to bind to the β5 site with greater selectivity compared to the human enzyme.

Furthermore, upon exploring the surfaces and binding interfaces of all the active sites in *Tv*20S and human 20S proteasomes, it becomes evident that the β2 and β5 active sites, along with their respective S1 and S3 pockets, exhibit significant differences in shape. These differences offer additional space for further inhibitor design (**Fig. 5**). Although the distinct amino acid composition and slight differences arrangement of the β2 compared to β5 active sites pose challenges for the design of selective inhibitors.

**Fig. 5:**
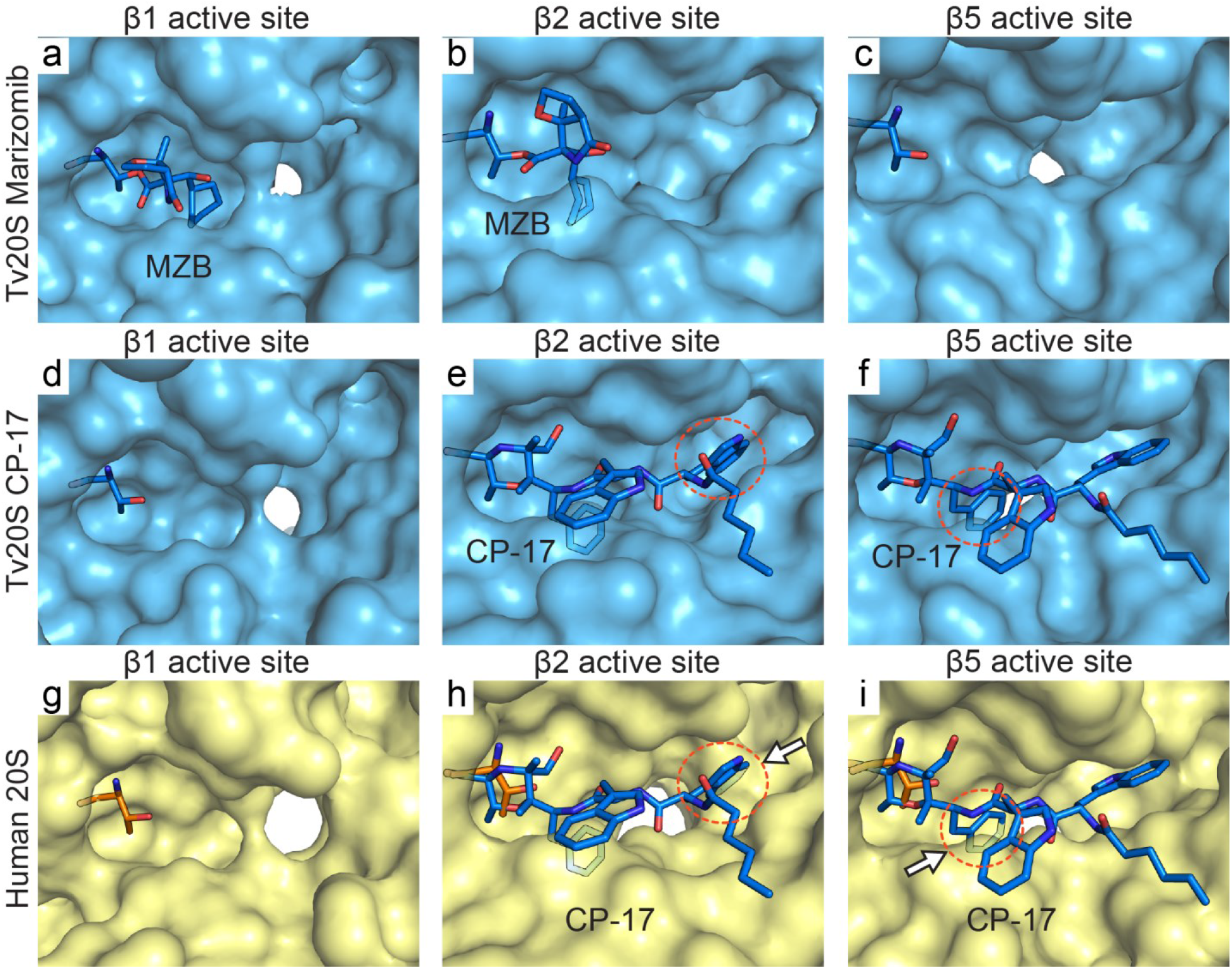
Surface detail of comparison of *Tv*20S and human 20S active sites. *Tv*20S surface is in light blue. Human 20S proteasomes is in light yellow and its Thr1 is shown as orange sticks. MZB and CP-17 are shown as blue sticks is displayed in human active site pockets using structural overlay from Fig.4. The clashes are encircled with a red dotted line. In panel h, Met130 of human β2 forms shallower binding pocket in this region whist similar elevation of active site pocket is observed formed by Met45 from human β5. In this location human β5 Met45 in this ligand-free structure is pushed by Ile35 towards the active site.

A comparison between the active sites of human and Trichomonas 20S proteasomes provides insights that may explain the differences in substrate specificity and lower cytotoxicity of CP-17, as previously reported ^8^. These disparities between human and Trichomonas proteasomes, along with unexplored areas, present potential opportunities for improved design strategies targeting the *Tv*20S proteasome.

## Discussion

Several drugs that target the proteasome have been approved for treatment of cancer and one proteasome inhibitor is in Phase II clinical trials for Leishmania. To develop proteasome inhibitors that are selective for a parasite proteasome over the human proteasome, structural and enzymatic studies are essential to uncover the subtle differences between these evolutionarily conserved complexes. However, the use of parasite proteasomes for research purposes has been limited by the availability of enzymes from native purification studies. This is particularly challenging for intracellular parasites such as *Plasmodium* and *Babesia* as they are difficult to cultivate in large quantities and require fresh human red blood cells. For other parasites such as *Trypanosoma, Leishmania* and *Trichomonas* species, isolation of enzyme with sufficiently high purity has been difficult. Recently, we have shown that native *Tv*20S co-purifies with actinin^8,22^. Numerous attempts to separate actinin from *Tv*20S failed and therefore an alternative approach was to express the recombinant enzyme complex. The human constitutive proteasome has been successfully expressed in baculovirus^24^ and therefore the first goal of our study was to make a recombinant parasite proteasome in the same system. We chose *Tv*20S as our model enzyme for these studies, as we have recently performed an in-depth substrate specificity profile of the three catalytic subunits and therefore have tools available to biochemically compare the native and recombinant enzymes^22^. As our long-term goal is to develop proteasome inhibitors of native *Tv*20S, we believe that it is important to ensure that the r*Tv*20S enzyme interacts with inhibitors in the same way as the native enzyme. This will avoid downstream misinterpretation of the structure-activity relationship. Successful expression of *Tv*20S proteasome in baculovirus expression system using an approach similar to what was performed for the human 20S proteasome^24^ demonstrates that this expression system could be transferable to proteasomes from other infectious organisms. In particular, we are interested in targeting the proteasome of the intracellular parasites, *Plasmodium falciparum* and *Babesia divergens*^32,33,52^ and the nematode *Schistosoma mansoni*^37^ as a lack of purified enzyme has hampered those studies.

Here, we present the two structures of the recombinant *Tv*20S proteasome bound to two different inhibitors. The overall architecture of *Tv*20S is conserved from archaea to humans. Four heptameric rings are stacked on top of each other, with the outer rings composed of the scaffolding α subunits and the inner rings composed of the catalytic β subunits. In archaea, each subunit is identical. However, in eukaryotes, the subunits have diverged, although they still exhibit high homology (**Fig. S3**). We were able to identify each *Tv20S* α and β subunit by comparing their sequences to the human homologs. We also identified a homolog of human Ump-1 in the *Tv* genome. Three distinct baculovirus containing the genes for 7α, 7β and *Tv*Ump-1 were used to infect Sf9 insect cells resulting in the production of fully folded recombinant *Tv*20S. All three catalytic subunits were active and they cleave the previously developed substrates. This is the first report of a pathogen proteasome being expressed in an insect cell line. Using CP-17, we showed conclusively that the β2 and β5 subunits of native and recombinant *Tv*20S are inhibited while also showing that β1 activity is increased in the presence of CP-17. Taken together, we are confident that the recombinant enzyme represents an ideal surrogate for future drug development.

Data from our structural studies reveal how MZB and CP-17 inhibit *Tv*20S and provide a structural basis for development of more specific inhibitors. For instance, the arrangement of the S2 substrate binding pocket in the β1 subunit prevents interactions with CP-17. This binding pocket contains Pro27, Gln112 and Gln127. The Gln residues are responsible for narrowing the binding grove which prevents binding of the P2 indole ring of CP-17. The active site pocket that accommodates the cyclohexenyl group of MZB appears to be too shallow to accommodate the phenyl group of CP-17. In the overlay of these β1 and β2 subunits, residue Arg45 of the β1 pocket is too close to phenyl group (2.4 A) (**Fig. S6, S7**)

Inhibitory measurements of enzyme activity demonstrated that MZB inhibits all three active subunits of the proteasome (β5, β2, and β1), thus targeting all six active sites. Consequently, MZB targets all six active sites present within the proteasome structure. In contrast, CP-17 selectively inhibits β5 and β2 subunits, therefore targeting four out of the six active sites. Our enzyme kinetic studies revealed that inhibition of β5 and β2 with CP-17 yielded an increase in substrate cleavage by the β1 subunit. To understand why catalytic activity increases when the other two subunits are inhibited, we compared the structure of the β1 subunit bound to MZB with that of the empty β1 subunit bound of the CP-17 structure. We anticipated that the empty β1 subunit may have a conformational change that allows easier access to the Ac-RYFD-amc substrate. However, the shape and volume of the empty and MZB-bound β1 subunits were identical (**Fig. 5a and 5d**) indicating that the increase in activity was not due to a structural change. We therefore predict that this increase in activity is due to the β1 substrate being directed towards the β1 subunit since the β2 and β5 subunits are blocked with CP-17. Therefore, its effective concentration at the β1 subunit is higher in the CP-17-treated enzyme than in the non-inhibited enzyme resulting in a 2-fold increase in activity.

Enzyme inhibition studies showed that CP-17 has higher selective for *Tv* over HeLa cells proves to be superior due to its index. This finding was confirmed in experiment measuring EC_50_ on and parasites. On the other hand, MZB exhibits a poorer selective index but demonstrates a better *in vitro* IC_50_ (**Fig. S8**).

CP-17 exhibits promising potential as a candidate for targeting the active sites of the *Tv*20S proteasome, demonstrating notable potency and specificity. This makes it an appealing hit molecule for structure-based design of a drug. CP-17 does not bind to the β1 site of Tv20S or c20S and an ideal inhibitor would bind to this site in Tv20S and not in c20S. Future analogues of CP-17 could leverage the presence of Cys46 in the S1 pocket of β1 by forming a covalent link with this residue, while still maintaining or even enhancing specificity for other binding sites. Significantly, since the human 20S proteasome lacks Cys46, direct targeting of Cys46 could greatly enhance the specificity of these subsequent compounds (**Fig. S6**). This would lead to improved inhibition of *Tv*20S proteasome activity by CP-17 derivatives, thereby further enhancing its therapeutic potential for the treatment of trichomoniasis.

## Acknowledgment

The cryo-EM data were collected at the UmeÅ Centre for Electron Microscopy (UCEM) and we thank in particular to Michael Hall, Camilla Holmlund and Linda Sandblad. We are grateful to Karim Rafie and Lar-Anders Carlson for invaluable help with cryo-EM experiments and data processing. The research was supported by NIH awards R01AI158612 and R21AI146387 to AJO, LE, and WHG. PF received funding from the European Union’s Horizon 2020 research and innovation program under the Marie Skłodowska-Curie grant agreement No [846688]. This research was funded by the project the National Institute Virology and Bacteriology (Programme EXCELES, Project No. LX22NPO5103) - Funded by the European Union - Next Generation EU (awarded E.B.). Academy of Sciences of the Czech Republic, RVO: 61388963, is also acknowledged. BMH was supported in part by the UCSD Graduate Training Program in Cellular and Molecular Pharmacology through an institutional training grant from the National Institute of General Medical Sciences, T32 GM007752. PF would like to acknowledge Milan Fabry for providing advice on cloning and Martin Horn for assistance with enzymatic assays. JA would like to acknowledge the St. Baldrick’s Foundation for the International Scholar award 2022–2025 and the deanship of scientific research at the University of Jordan for the scientific leave. We would like to thank Rowan O’Donoghue for artwork.

## Author Contribution

J.S. collected and processed cryo-EM data, P.F., B.M.H performed experiments. Y.M. and L.E cultured *T. vaginalis* and provided cell lysate, J.A and W.H.G designed and synthesized CP-17, P.F., A.J.O. and E.B conceived the project. J.S., P.F., A.J.O and E.B. wrote the manuscript. A.J.O. and E.B. supervised the project. P.F., L.E., W.H.G., A.J.O., and E.B. obtained funding.

## Data availability

The atomic coordinates and cryo-EM density maps were deposited in the Protein Data Bank (https://www.rcsb.org) under the PDB accession codes 8OIX (EMD-16901), 8P0T (EMD-17337).

