## Supplementary Information for "Structural elucidation of recombinant *Trichomonas vaginalis* 20S proteasome bound to covalent inhibitors"

Fig. S1**. Cloning plasmids and insert sequences.**

Schematic representation of the three cloning plasmids constructed from the original plasmid (denoted as "pACEBac1") and the insert (BamHI_α1_α2_α3_α4_α5_α6_α7_NotI, BamHI_β1_β2_β3_β4_β5_β6_β7+tag_NotI and BamHI_Ump-1_NotI). Each plasmid is labeled as α, β, or Ump-1, respectively. The cloning process involved the insertion of specific fragments into the original plasmid backbone. The figure was created using SnapGene® software (from Dotmatics; available at snapgene.com). Nucleotide sequence of the inserted DNA used in the cloning process are below. The sequence is shown in the 5' to 3' direction. The restriction sites are highlighted.

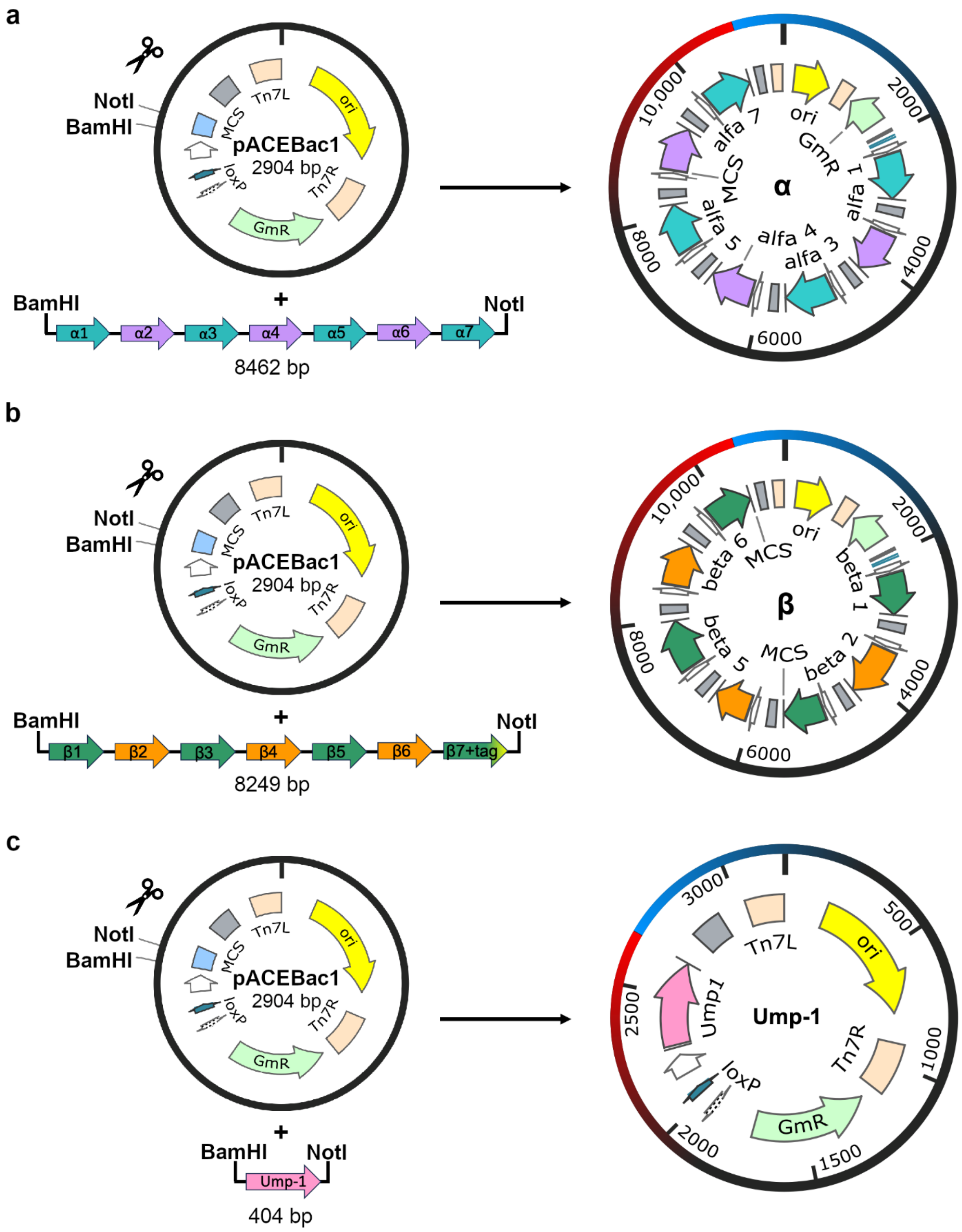

Insert BamHI_α1_α2_α3_α4_α5_α6_α7_NotI

```
ggatccGCCACCATGAGCAGCGGTGCAGATCGTTATCTGACCGTTTTTAGTGCCGAAGGTCGTCTGTGGCAGGTTGAATATAGTTTTAA
AGCAGTTAAACAGGCCGAAGTTACCGCAGTTGCAGTTAAAAGCAAAAATGCAGTTTGTGTTGCCGTGCAGAAAAAGGTTAGCGATAAAC
TGATTGATCCGAGCACCGTTACACACATGTATCGTATTACCGATAATGTTGGTGCATGTCTGGTTGGTCTGCCGAGTGATGTTAACTTT
ATTGTTATGCTGCTGCGTAGCTTTGCCAACAACTTTGAATATAAACAGGGCTTTAGCATCCCGGTTTCAATTCTGGCACAGATGCTGAG
CGAACGTCATCAGCTGGAAAGCCAGCTGGTTTATGTTCGTCCGAGCGCAGTTAGCGCAATTCTGTTTGGTCTGGATGGTCCGAGCGATA
GCTTTGCACTGTATAAAATCGAACCGAGCGGTTATAGCAATGGTTTTCGTGCAGTTGCATGTGGCGTTAAAGAAATTGAAGCAATGAGC
GCACTGGAAAAGAAGATGGAAGATTTTGAAACACCGGAAGCAACCGCAGAATTTACCCTGAGCACCCTGCAAACCGTTTGTGGTGTTGA
TTTTGAAGCACAGGATGTTGAAGTTAGCCTGCTGACCCGTGATAATAGCAAATTTTCAAAACTGCCGAACGACAAGGTGAACGAAATTC
TGCATGCCGTTGCCGAAAAAGATTAAAGATCTAGAGGATCATAATCAGCCATACCACATTTGTAGAGGTTTTACTTGCTTTAAAAAACC
TCCCACACCTCCCCCTGAACCTGAAACATAAAATGAATGCAATTGTTGTTGTTAACTTGTTTATTGCAGCTTATAATGGTTACAAATAA
AGCAATAGCATCACAAATTTCACAAATAAAGCATTTTTTTCACTGCATTCTAGTTGTGGTTTGTCCAAACTCATCAATGTATCTTATCA
TGTCTGGATCTGATCACTGCTTGAGCCTAGAAGATCCGGCTGCTAACAAAGCCCGAAAGGAAGCTGAGTTGGCTGCTGCCACCGCTGAG
CAATAACTATCATAACCCCTAGGGTATACCCATCTAAGGTAGCGAGTTTAAACACTAGTATCGATTCGCGACCTACTCCGGAATATTAA
TAGATCATGGAGATAATTAAAATGATAACCATCTCGCAAATAAATAAGTATTTTACTGTTTTCGTAACAGTTTTGTAATAAAAAAACCT
ATAAATATTCCGGATTATTCATACCGTCCCACCATCGGGCGCGAGCTCGCCACCATGGGCGATAGCGATTTTAGCCTGACCACCTTTAG
CAGCGGTGGTAAACTGAATCAGATTGAAAGCGCACTGAAAGCAGTTAGCTTAGGTGGCCAGTGTGTTGGTGTTAAAGCCAAAAATGGTG
CAGTTATTGCCTGTGAAAGCAAACCGAGCAGTCCGCTGGTTGAAAAAGTTACCAATCTGAAAGTGCAGAAAATCAACGATAATGTGGGC
ATTGTTTATAGCGGTGTGAACACCGATTTTCACGTTATTCTGAAAAGCCTGCGTAAAGCCAGCATCAAATATAGCCTGCGTCTGGGTGT
TGAAATGCCGACACGTGAAGTTGTTAAACATGCAGCACATAAGATGCAGTATTATACCCAGATTGGTGGTGTTCGTCCGTTTGGTGTTA
GCCTGCTGATTATTGGTTGGGAAGAACTGGGTCCGACACTGTGGCAGGTTGATCCGAGCGGCACCTTTTGGGCATGGAAAGCAACCGCA
CTGGGTAAACGTAGTGATGGTAGCCGTACCTTTCTGGAACGTCGTTATAGCGAAGATCAGAGCGTTGATGATGCAATTCATACCGCAAT
TAGCACCCTGAAAGAAGGTTTTGACGGCCAGCTGACCGCAGAACTGATTGAAATTGGTGTTGTTGATGAAACCCGTAAATTTCGTACCC
TGAGCACCGCAGAAATCCGCGATTTTCTGACCGAAGTTTAAGAATTCAGAGGATCATAATCAGCCATACCACATTTGTAGAGGTTTTAC
TTGCTTTAAAAAACCTCCCACACCTCCCCCTGAACCTGAAACATAAAATGAATGCAATTGTTGTTGTTAACTTGTTTATTGCAGCTTAT
AATGGTTACAAATAAAGCAATAGCATCACAAATTTCACAAATAAAGCATTTTTTTCACTGCATTCTAGTTGTGGTTTGTCCAAACTCAT
CAATGTATCTTATCATGTCTGGATCTGATCACTGCTTGAGCCTAGAAGATCCGGCTGCTAACAAAGCCCGAAAGGAAGCTGAGTTGGCT
GCTGCCACCGCTGAGCAATAACTATCATAACCCCTAGGGTATACCCATCTAAGGTAGCGAGTTTAAACACTAGTATCGATTCGCGACCT
ACTCCGGAATATTAATAGATCATGGAGATAATTAAAATGATAACCATCTCGCAAATAAATAAGTATTTTACTGTTTTCGTAACAGTTTT
GTAATAAAAAAACCTATAAATATTCCGGATTATTCATACCGTCCCACCATCGGGCGCCCGCGGGCCACCATGACCTATCGTTATGATGC
AGGCACCACCACCTTTAGCAGTGATGGTCGTATTCTGCAGGTTGAATATGCAATTCAGAGCATTAATCAGGCAGGCACCGCAATTGGTG
TTCAGTTTACCAATGGTGTTGTTCTGGCAGCCGAAAAGAAAAATACCGGTCGTCTGGTTGATTACCTGTTTCCTGAGAAAATGGCCAAA
ATTGATGGTCATATTGTTACCGCAGTTGCAGGTCTGACCGCAGATGCAAATACCCTGGTTGATCTGATGCGTACCAGCGCACAGAAATA
TCTGAAAACCTATGATGAGCAGATGCCGGTTGAACAGCTGGTTCGTATGGTTTGTGATGAAAAACATAGCTATACCCAGTATGGTGGTC
TGCGTCCGTATGGTGTTAGCTTTCTGATTGCAGGTTATGATCGTCATAAAGGTTGTCAGCTGTATCTGACCGATCCGAGCGGTAATTTT
GGTGGTTGGAAAGCAACCGCCATTGGTGAAAATAATCAGACCGCACAGAGCATTCTGAAAAGCCAGTATAAAGATAATATGACCGCAAC
CGAAGCAATGGATCTGACCGTTAAAGTTCTGTGTAAAACCCTGGATAGCACCAGCCTGAGCGCAGATAAACTGGAATTTGCAGTTCTGC
AGTTTCGCGAAGAATATGGTCCGAAAGTGCGTATTCTGACCACCAGTGAAGTTGATACCCTGATGAAACGTTATGAGGAAACCATTAAA
AAGTCAGCCGAGGAAAAAGAATAAGGTACCAGAGGATCATAATCAGCCATACCACATTTGTAGAGGTTTTACTTGCTTTAAAAAACCTC
CCACACCTCCCCCTGAACCTGAAACATAAAATGAATGCAATTGTTGTTGTTAACTTGTTTATTGCAGCTTATAATGGTTACAAATAAAG
CAATAGCATCACAAATTTCACAAATAAAGCATTTTTTTCACTGCATTCTAGTTGTGGTTTGTCCAAACTCATCAATGTATCTTATCATG
TCTGGATCTGATCACTGCTTGAGCCTAGAAGATCCGGCTGCTAACAAAGCCCGAAAGGAAGCTGAGTTGGCTGCTGCCACCGCTGAGCA
ATAACTATCATAACCCCTAGGGTATACCCATCTAAGGTAGCGAGTTTAAACACTAGTATCGATTCGCGACCTACTCCGGAATATTAATA
GATCATGGAGATAATTAAAATGATAACCATCTCGCAAATAAATAAGTATTTTACTGTTTTCGTAACAGTTTTGTAATAAAAAAACCTAT
AAATATTCCGGATTATTCATACCGTCCCACCATCGGGCGCCTCGAGGCCACCATGAGCGATTATACCCGTAGCATTACCCGTTTTAGTC
CGGATGGTCGTCTGTTTCAGATTGATCATGCACATGCAGCAGTTCAGCGTGGCACCACCGTTGTTGCAACCCGTAGTAAAGATATGATT
GTTATTGCCGTTGAGAAAACCGCAGTTGCAAAACTGCAAGATCCGCATACCTTTAGCAAAATTTGTAGCCTGGATAAACATGTGATGTG
TGCATTTGCAGGTCTGCATGCCGATGCACGTCGCCTGATTCAGAGCGGTCAGCGTCAGTGTCAGAGCCATCGTCTGACCTATGAAGATC
CGATTAGCATTGAAAACATTGCCCGTTATATTGCAACCCTGCAACTGAAAAATACCCAGAGCGGTGGTGCACGTCCGTATGGTGTTAGC
ACCCTGATTTGTGGTTTTGATGATATGACCAGCCAGCCGCATATTTATGAAACCCTGCCGAGCGGCACCTATGCAGAATGGAAAGCACG
TACCATTGGTCGTCATGATCAGACCGTTATGGAATATCTGGAAAAACACTACAAAGACGATATGACCGATGAAGAAGCACAGAAACTGG
CAATTGGTGCACTGCTGGAAGTTGTTGAAAATGGTAGCAAAAATCTGGAAGTGGCCTATATGAAACGTGGTGGTACAATGGAAATTATG
GCCGAAGAGGTTCTGGATGCACTGATTGAAAGCACCAAAGCAAAATAAACGCGTAGAGGATCATAATCAGCCATACCACATTTGTAGAG
GTTTTACTTGCTTTAAAAAACCTCCCACACCTCCCCCTGAACCTGAAACATAAAATGAATGCAATTGTTGTTGTTAACTTGTTTATTGC
AGCTTATAATGGTTACAAATAAAGCAATAGCATCACAAATTTCACAAATAAAGCATTTTTTTCACTGCATTCTAGTTGTGGTTTGTCCA
AACTCATCAATGTATCTTATCATGTCTGGATCTGATCACTGCTTGAGCCTAGAAGATCCGGCTGCTAACAAAGCCCGAAAGGAAGCTGA
GTTGGCTGCTGCCACCGCTGAGCAATAACTATCATAACCCCTAGGGTATACCCATCTAAGGTAGCGAGTTTAAACACTAGTATCGATTC
GCGACCTACTCCGGAATATTAATAGATCATGGAGATAATTAAAATGATAACCATCTCGCAAATAAATAAGTATTTTACTGTTTTCGTAA
CAGTTTTGTAATAAAAAAACCTATAAATATTCCGGATTATTCATACCGTCCCACCATCGGGCGCAAGCTTGCCACCATGTTTAATAGCG
GCAGCGAATATGATCGCAATGTGAATACCTTTAGTCCGGATGGTCGTCTGCTGCAAGTTGAATATGCAATTGAAGCAGTTAAACTGGGT
AGCAGCGCAGTTGCAATTCTGTGTCCGGAAGGTGTTATTTTTGCAGTTGAAAAACGTCTGAGCAGCCAGCTGCTGATTGCAAGCAGCGT
TGAAAAAGTTTATGCCATCGATGATCATGTTGGTGTTGTTATGGCAGGTCTGGCAGCAGATGGTCGTACAATGGTTGAACACATGCGTG
TTGAAGCACAGAATCATCGTTTTAGCTTTGATGAACCGATTGGCATTAAAGCAGTTACCCAGAGCGTTTGTGATCTGGCACTGGCATTT
GGTGAAGGTCGTCGTAAGAAAGGTGATGGTCAGATGAGCCGTCCGTTTGGCACCGCACTGCTGGTTGCAGGTATTGAAAATGGTAAATG
TCACCTGTTTCATACCGATCCGAGCGGCACCTATACCGAATGTCGTGCACGTGCCATTGGTGGTGGTAGCGAAGGTGCCGAAGCACTGC
TGCGTGATCTGTATAAAGATGGTATGACCCTGCATGAAGCAGAGGATCTGGCCCTGAGCACCCTGCGTCAGGTTATTCAAGAAAAACTG
AATGAGAACAATGTGGAAGTTGCATGTGCACGTGTTAGCACCGGTAAATTTGAAATCTATACCAGCGAACAGCGCCAAGAAATTGTTGC
CCGTCTGCCTCCGCCTATTATTCCGGAATAAGTCGACAGAGGATCATAATCAGCCATACCACATTTGTAGAGGTTTTACTTGCTTTAAA
```

AAACCTCCCACACCTCCCCCTGAACCTGAAACATAAAATGAATGCAATTGTTGTTGTTAACTTGTTTATTGCAGCTTATAATGGTTACA
AATAAAGCAATAGCATCACAAATTTCACAAATAAAGCATTTTTTTCACTGCATTCTAGTTGTGGTTTGTCCAAACTCATCAATGTATCT
TATCATGTCTGGATCTGATCACTGCTTGAGCCTAGAAGATCCGGCTGCTAACAAAGCCCGAAAGGAAGCTGAGTTGGCTGCTGCCACCG
CTGAGCAATAACTATCATAACCCCTAGGGTATACCCATCTAAGGTAGCGAGTTTAAACACTAGTATCGATTCGCGACCTACTCCGGAAT
ATTAATAGATCATGGAGATAATTAAAATGATAACCATCTCGCAAATAAATAAGTATTTTACTGTTTTCGTAACAGTTTTGTAATAAAAA
AACCTATAAATATTCCGGATTATTCATACCGTCCCACCATCGGGCGCCCCGGGGCCACCATGTTCCGCAGCAAATATGATGAAAATGCC
ACCACCTTTAGTCCGGAAGGTCGTATTCTGCAAGTTGAAAATGCAATGAAAGCAGTTCAGCAGGGTATGCCGACCGTTGGTCTGAAAAG
CAAAACCCATGCAGTTATTGCCGGTGTTATGCATAGCCCGAGCGAATTTAGCAGCCATCAGCCGAAAATCTTTAAAATCGATCAGCATA
TTGGTGTGGCAATTAGCGGTCTGACCGCAGATGGTCGTGGTCTGTGTAAATTTCTGCGTAATGAATGTCTGCATCACACCTTTTGTTTT
GGCACCGAAATTCGTGTTGCCGATCTGGCAGATACCGTTGCACTGCAAAGCCAGAAAAAGACCAGCAAAGTTGGTAAACGTCCGTATGG
TGTTGGTCTGCTGATGATTGGTGCGGGTGTTGATGGTCCGCGTCTGTTTGAAACCTGTCCGAGCGGTCAGCATTGGGAATATAATGCAC
AGGCAATTGGTCGTCGTGCCCAGGCAGCAAAAACCTATCTGGAAACCAATCTGAATGAATTTCCGGATTGTACCCGTGATCAACTGATT
CGTCATGCACTGCGTGCACTGAATGATTGTAAAAGCCGTGAAAGCGATAGCCTGGAAGCAATTGCACTGGGTGTTGTTGGTATTGATGA
ACCGTTTACCATTCTGGAAGGTCCGGAATTACAGAAATATATCGATTAAAGGCCTAGAGGATCATAATCAGCCATACCACATTTGTAGA
GGTTTTACTTGCTTTAAAAAACCTCCCACACCTCCCCCTGAACCTGAAACATAAAATGAATGCAATTGTTGTTGTTAACTTGTTTATTG
CAGCTTATAATGGTTACAAATAAAGCAATAGCATCACAAATTTCACAAATAAAGCATTTTTTTCACTGCATTCTAGTTGTGGTTTGTCC
AAACTCATCAATGTATCTTATCATGTCTGGATCTGATCACTGCTTGAGCCTAGAAGATCCGGCTGCTAACAAAGCCCGAAAGGAAGCTG
AGTTGGCTGCTGCCACCGCTGAGCAATAACTATCATAACCCCTAGGGTATACCCATCTAAGGTAGCGAGTTTAAACACTAGTATCGATT
CGCGACCTACTCCGGAATATTAATAGATCATGGAGATAATTAAAATGATAACCATCTCGCAAATAAATAAGTATTTTACTGTTTTCGTA
ACAGTTTTGTAATAAAAAAACCTATAAATATTCCGGATTATTCATACCGTCCCACCATCGGGCGCCATATGGCCACCATGAGCGGTGCA
GGTAGCGGTTATGATTTTAATCCGATTACCTTTAGTCCGGATGGTCGTCAGTTTCAGGTTGAATATGCAACCAAAGCCGTTGAAAAAGA
TAGCCTGGCACTGGGTGTTAAATGCAAAGATGGTATTCTGCTGGCAGCCGAGAAAAATCTGACCAGCACACTGCTGACCCCTGGTGGTA
ATCCGCGTATCTTTTGGATTAATGATAGCATTGCATGTGCCACCATTGGTCATCGTCCGGATTGTTATAGCATTGTTGAACAGAGCCGT
AATCGTGCAGAAACCTTTACCAGCAATTTTGGCATCAAAATTACCGTTCCGCAGCTGGCAAGCGAAGTTAGCCAGCAGTTTCATCTGGC
ACATTATTATCAGGCATATCGTCCGTTTGGTTGTACCGTTATTTTTGCCAGCTATAAAGATGATGCCCTGTATGCAATTGAACCGAGCG
GTGCCTTTTATGGTTATTTTGCAAGCTGCTTTGGCAAGAATAGCAATCTGGCACGTGCAGAATTACAGAAAACCGAATGGAAAAATATC
ACCGTTCGTGAAGCAGTTCCGGAAGTTGCACGTATTATCAAAAGCCTGCATGAAAGCCAGTTTAAAAAGTGGGAAATCGAAATGTTTTG
GCTGTGCGAAGAAACCAATGGTCGTCCGCAGAAAGTGCCGGAAGATGTTTTTCAGAGCCGTTTTGTTAATGAAAACCCGCAGAATTAAg
cggccgc

Insert BamHI_β1_β2_β3_β4_β5_β6_β7+tag_NotI

*ggatcc*GCCACCATGAGCGAATATCAGTTTCCGAAAGAAAGCATGGGTACAACCCTGCTGGCAATTCAGTGTACCGATGGTGTTGTTAT
GGCAAGCGATAGCCGTACCAGCAGCGGTAGCTTTATTCCGAATCGTGCAACCAACAAAATTACCGAAATTCAGCCGAAAATCTTTGCAG
CACGTTGTGGTAATGCAGCAGATACCCAGTTTCTGGCACGTGCAGTTAAAAACTATCTGAATGCACTGAACATCACCCGTGAAAATACC
GATGATAGCACCATTCTGGTTGCCAGCAATGTTATTCGTAGCCTGATTGTTCGTTATCGTCAGTATCTGAGTGCCGGTGTTATTGTTGG
TGGTTGGGATAGCGCAGGTCCGCAGGTTTATAGCATTGAAGTTAGCGGTATGGCCATCAAAAAGAAAATTGCAAGCAATGGTAGCGGCA
GCACCTATATTCAGGCATATATTGATCAGAACTATCGCGAGGATATGACAATGGAAGAAGCAACCAAATTTGCAATTGCCGCAGTTACC
GGTGCAATTATTCGTGATGGTAGCAGCGGTGGTGTTGTGAATATTGTTCAGATTAATGCAGATGGTGCCAAACGTATGACCGTTCGTCC
GGCACAGCAGCCGTTTAACTATGATATTGTTAAAGGTTAAAGATCTAGAGGATCATAATCAGCCATACCACATTTGTAGAGGTTTTACT
TGCTTTAAAAAACCTCCCACACCTCCCCCTGAACCTGAAACATAAAATGAATGCAATTGTTGTTGTTAACTTGTTTATTGCAGCTTATA
ATGGTTACAAATAAAGCAATAGCATCACAAATTTCACAAATAAAGCATTTTTTTCACTGCATTCTAGTTGTGGTTTGTCCAAACTCATC
AATGTATCTTATCATGTCTGGATCTGATCACTGCTTGAGCCTAGAAGATCCGGCTGCTAACAAAGCCCGAAAGGAAGCTGAGTTGGCTG
CTGCCACCGCTGAGCAATAACTATCATAACCCCTAGGGTATACCCATCTAAGGTAGCGAGTTTAAACACTAGTATCGATTCGCGACCTA
CTCCGGAATATTAATAGATCATGGAGATAATTAAAATGATAACCATCTCGCAAATAAATAAGTATTTTACTGTTTTCGTAACAGTTTTG
TAATAAAAAAACCTATAAATATTCCGGATTATTCATACCGTCCCACCATCGGGCGCGAGCTCGCCACCATGGAAGCAGGTCTGGGTTTT
GATTTTAGCAATTATGCCCGTAACAAAAGCCTGGAACCGAAACTGGGTAAACCGCAGCTGCTGAGCACCGGCACCACCATTGCAGCAGC
AATTTTTGATGGTGGTGTTGTTCTGGGTGCAGATACCCGTGCAACCGCAGGTCCGATTGTTGCAGTTAAAGATGAAATGAAACTGCACT
ACATCAGCGATAACATTTGGGTTTGTGGTGCAGGTATTGCAGCCGATAATGATAACATTAATGCAGTGATTAGCGCCAAACTGCGTCTG
TTTCAGATGAATACCGGTCTGCAACCGCGTGTTGATCAGTGTACCAATATTCTGGCAAGTCGCCTGTTTCAGTATATGGGTTATATTCA
GGCAGCACTGATTGTTGGTGGTATTGATTTTCAGGGTCCGCAGGTTTATCAGGTTGCACCGCATGGTAGCTTTAGCAAACAGCCGTTTA
TTGCACAAGGTAGCGGTAGCCTGGCAGCAATTAGCGTTCTGGAAAATCGTTGGCATAACAAAATGAACGAACACGACTGTATGGAAATG
GTTGCAGATGCAATTTATGCCGGTATTACCAATGATTTAGGTTCAGGTAGCCATGTTAATCTGTGCGTGATTAAACGTGAAAACCCGGA
AGATAAACAGAGCAAAGTGATCTACACCTTCTATAAAGATTATCGCGTGCCGCATGAAAACGATCGTAATTTTCGTCTGGAACCGCAGA
TCAATAACATTGATGTGGAAGTGATTAAAACCACCGAACGTCCGCTGACACTGCCGGATGTTCATCTGGAAATTCTGGATGATGCACCG
GCATAAGAATTCAGAGGATCATAATCAGCCATACCACATTTGTAGAGGTTTTACTTGCTTTAAAAAACCTCCCACACCTCCCCCTGAAC
CTGAAACATAAAATGAATGCAATTGTTGTTGTTAACTTGTTTATTGCAGCTTATAATGGTTACAAATAAAGCAATAGCATCACAAATTT
CACAAATAAAGCATTTTTTTCACTGCATTCTAGTTGTGGTTTGTCCAAACTCATCAATGTATCTTATCATGTCTGGATCTGATCACTGC
TTGAGCCTAGAAGATCCGGCTGCTAACAAAGCCCGAAAGGAAGCTGAGTTGGCTGCTGCCACCGCTGAGCAATAACTATCATAACCCCT
AGGGTATACCCATCTAAGGTAGCGAGTTTAAACACTAGTATCGATTCGCGACCTACTCCGGAATATTAATAGATCATGGAGATAATTAA
AATGATAACCATCTCGCAAATAAATAAGTATTTTACTGTTTTCGTAACAGTTTTGTAATAAAAAAACCTATAAATATTCCGGATTATTC
ATACCGTCCCACCATCGGGCGCCCGCGGGCCACCATGAGCGATATCAGCACCTATAATGGTAGCTGTGTTCTGGCAATGGCAGGCGATC
ATTGTGTTGCAATTGCCAGCGATCGTCGTCTGGGTGTTAATATGCTGACCGTTAGCAAAGATTTCAAACGCATCTTTCAGATCAACGAC
CGCATTTATCTGGGTTTAGCAGGTCTGGCAACCGATGTTCTGACCGTTCGTGAACAGCTGCGTTTTGATGTTAATCTGCTGGAACTGCG
TGAAGAACGTCCGATTGATCCGAAAAAGTTTATGAATCTGGTTAAAAGCACCCTGTACGAAAAACGCTTTAGCCCGTTCTTTGTTACAC
CGGTTATTGCAGGTCTGCTGCCGGAAACCAATGAACCGTATCTGGCAGCAAGCGATAGCATTGGTGCATTTGCATTTCCGAAAGATTTT

GCAGTTGCAGGCACCTGTGAAGAAAGCCTGTATGGTATTTGTGAAAGCGCATGGCGTCCGAATATGAATCCGGATGAACTGTTTGAATG
TACCGCCAAATGTCTGATTGCAGCAGTTGAACGTGATAGCATTAGCGGTTGGGGTGGTATTGTTTATATCATCACCCAGGATAAAGTGA
TCATCAAAGAAATCAAAACCCGCATGGATTAAGGTACCAGAGGATCATAATCAGCCATACCACATTTGTAGAGGTTTTACTTGCTTTAA
AAAACCTCCCACACCTCCCCCTGAACCTGAAACATAAAATGAATGCAATTGTTGTTGTTAACTTGTTTATTGCAGCTTATAATGGTTAC
AAATAAAGCAATAGCATCACAAATTTCACAAATAAAGCATTTTTTTCACTGCATTCTAGTTGTGGTTTGTCCAAACTCATCAATGTATC
TTATCATGTCTGGATCTGATCACTGCTTGAGCCTAGAAGATCCGGCTGCTAACAAAGCCCGAAAGGAAGCTGAGTTGGCTGCTGCCACC
GCTGAGCAATAACTATCATAACCCCTAGGGTATACCCATCTAAGGTAGCGAGTTTAAACACTAGTATCGATTCGCGACCTACTCCGGAA
TATTAATAGATCATGGAGATAATTAAAATGATAACCATCTCGCAAATAAATAAGTATTTTACTGTTTTCGTAACAGTTTTGTAATAAAA
AAACCTATAAATATTCCGGATTATTCATACCGTCCCACCATCGGGCGCCTCGAGGCCACCATGCTGAGCATTGTTGGTCTGCAAGGTCC
GGATTGGGTTCTGATTGCAGCAGATAGCAGCGTTAGCAGCAGCATTATTTGTATGAGCGAAAACTATGATCGTATCGCACAGCTGGATG
ATCGTCATGCACTGGCAATGAGCGGTGAAACCGGTGATTGTCTGCAACTGAGCGAATATTTACAGGGTAATGTTGCCCTGTACAAATTT
CGTAATGGTGTTGAACTGAGCAGTGATGCCCTGGCACATTTTATTCGTCATACAATGGCAAAAGCCGTTCGTAAAAGCCCGTATGAAGT
TAATATGCTGCTGAGCGGTTATGATGGTAAACCGCATCTGTATTTCATGGATTATCTGGGCACCCTGCAAAGCATTCCGTATGGTGCAC
AGGGTTATTGTCAGTATTTTGTGATGAGCGTGTTCGATAAGCACTATAAAGAAGGTCTGACCCTGGAAGATGGTAAAGAACTGATGAAA
CTGGCACTGAACCAGATTAAACAGCGTTTTACCGTTGCACCGCATGGCTTTATTGTTAAACTGGTGGATAAAAACGGCATCACCAAAAT
CGATCTGGAATAAACGCGTAGAGGATCATAATCAGCCATACCACATTTGTAGAGGTTTTACTTGCTTTAAAAAACCTCCCACACCTCCC
CCTGAACCTGAAACATAAAATGAATGCAATTGTTGTTGTTAACTTGTTTATTGCAGCTTATAATGGTTACAAATAAAGCAATAGCATCA
CAAATTTCACAAATAAAGCATTTTTTTCACTGCATTCTAGTTGTGGTTTGTCCAAACTCATCAATGTATCTTATCATGTCTGGATCTGA
TCACTGCTTGAGCCTAGAAGATCCGGCTGCTAACAAAGCCCGAAAGGAAGCTGAGTTGGCTGCTGCCACCGCTGAGCAATAACTATCAT
AACCCCTAGGGTATACCCATCTAAGGTAGCGAGTTTAAACACTAGTATCGATTCGCGACCTACTCCGGAATATTAATAGATCATGGAGA
TAATTAAAATGATAACCATCTCGCAAATAAATAAGTATTTTACTGTTTTCGTAACAGTTTTGTAATAAAAAAACCTATAAATATTCCGG
ATTATTCATACCGTCCCACCATCGGGCGCAAGCTTGCCACCATGCAGAGCCTGTATCTGAAACCGCGTGATGAAGTTGAAAGCGAAGAA
ACCACCAAAGCACTGGAACCGGCACATGTTCCAGATCCGTGTCAGTTTGTTAAAAACCATATTAGCCTGAGCTATACCAACGAACCGGG
TAAAATGGCAGCAGTTCATGGCACCACCACACTGAGCTTTATCTATAATGGTGGTATTGTTGTTGCCGTTGATAGCCGTGCAACCGGTG
GTCAGTTTATCTTTAGCCAGACCGTTATGAAAATTCTGCCGCTGGCACCGAATATGATTGGTACAATGGCAGGCGGTGCAGCAGATTGT
CAGTATTGGCTGCGTAATCTGAGCCGTCTGATTCAGCTGCATAAATTTCGTTATCAGCAGCCGCTGACCGTTGCAGCAGCAAGCAAAAT
TCTGGTTAATGAACTGTATCGCTACAAGGGCTATAATCTGAGTATTGGTAGCATGATTTGCGGCTATGATAATACCGGTCCGCACATCT
TTTATATCGATAATCATGGTAGCCGCATTGCCGGTAAACGTTTTAGCGTTGGTAGCGGTAGCACCCATGCCTATGGTGTTCTGGATACC
TGTTATCGTGAAGATATGACCAAAGAAGAAGCCTGCGAACTGGGTCGTCGTGCCATTTATCATGCAACCTATCGTGATAGCGGTAGTGG
TGGTCGTGTTAGCGTTGTTCATATTACCCAGAATGGTGTGGAATGGATCGATAAAACCGATGTGTTTGATATGCACGACTTTAGCAAAA
CCACCTTTTAAGTCGACAGAGGATCATAATCAGCCATACCACATTTGTAGAGGTTTTACTTGCTTTAAAAAACCTCCCACACCTCCCCC
TGAACCTGAAACATAAAATGAATGCAATTGTTGTTGTTAACTTGTTTATTGCAGCTTATAATGGTTACAAATAAAGCAATAGCATCACA
AATTTCACAAATAAAGCATTTTTTTCACTGCATTCTAGTTGTGGTTTGTCCAAACTCATCAATGTATCTTATCATGTCTGGATCTGATC
ACTGCTTGAGCCTAGAAGATCCGGCTGCTAACAAAGCCCGAAAGGAAGCTGAGTTGGCTGCTGCCACCGCTGAGCAATAACTATCATAA
CCCCTAGGGTATACCCATCTAAGGTAGCGAGTTTAAACACTAGTATCGATTCGCGACCTACTCCGGAATATTAATAGATCATGGAGATA
ATTAAAATGATAACCATCTCGCAAATAAATAAGTATTTTACTGTTTTCGTAACAGTTTTGTAATAAAAAAACCTATAAATATTCCGGAT
TATTCATACCGTCCCACCATCGGGCGCCCCGGGGCCACCATGGAAGGTGAATTTCGCGAGAATAAGAAAGGTCAGTGGTCACCGTATGA
AATGCATGGTGGCACCGCAATTGGTATTTGTGGTGATGATTATGTTGTGATTGGTGCAGATACCCGTCTGAGCGTTGATTATAGCATTG
ATAGCCGTCATAAAGCCCGTATCTTTAAGATGAATAGCAACTGTATGATTAGCGCCACCGGTTTTGATGGTGATATTGATGCATTTATT
ACCCGCATGCGTAGCATTCTGCTGAATTATGAAAACCAGCACTTTCACGAAATGAGCGTGGAAAGCGTTGCACGTTGTGTTAGCAATAC
CCTGTATAGCAAACGTTTCTTCCCGTACTATATCAACATTCTGGTTGGTGGCATTAACAGCGAAGGTAAAGGTAAACTGTATGGTTATG
ATCCGGTTGGCACCATTGAGGATCTGCATTATGATAGCAATGGTAGCGGTAGCAGCCTGGCAGCACCGCTGCTGGATAGCGCATTTGGT
ACAATTCATCATAATACCCGTCCGTTTCCGGCAGTTTCACTGCAAGATGCCAAAAACATTGTTCGTGATGCAATTTGTAGCGTTACCGA
ACGTGATATCTATACCGGTGATGCACTGCAACTGTGTGTTTTTACCAAAGATGGTTTTGCCCAAGAAGAATTTCCGCTGCCTCGTCATT
AAAGGCCTAGAGGATCATAATCAGCCATACCACATTTGTAGAGGTTTTACTTGCTTTAAAAAACCTCCCACACCTCCCCCTGAACCTGA
AACATAAAATGAATGCAATTGTTGTTGTTAACTTGTTTATTGCAGCTTATAATGGTTACAAATAAAGCAATAGCATCACAAATTTCACA
AATAAAGCATTTTTTTCACTGCATTCTAGTTGTGGTTTGTCCAAACTCATCAATGTATCTTATCATGTCTGGATCTGATCACTGCTTGA
GCCTAGAAGATCCGGCTGCTAACAAAGCCCGAAAGGAAGCTGAGTTGGCTGCTGCCACCGCTGAGCAATAACTATCATAACCCCTAGGG
TATACCCATCTAAGGTAGCGAGTTTAAACACTAGTATCGATTCGCGACCTACTCCGGAATATTAATAGATCATGGAGATAATTAAAATG
ATAACCATCTCGCAAATAAATAAGTATTTTACTGTTTTCGTAACAGTTTTGTAATAAAAAAACCTATAAATATTCCGGATTATTCATAC
CGTCCCACCATCGGGCGCCATATGGCCACCATGCAGGTTATTACCGCAAGCGGTGCAATTGTTGCAGCAAAATATGATGGTGGTATTCT
GCTGGCAAGCGATCTGAGCATTACCTATGGTAGCATGTTTCGCCATAATAACGTTAGCCATTTTGTTGAAGTTGCACCGAACATTATTA
TCGGTGCCAGCGGTGAATTTGCAGATTTTCAGACCCTGATTGAAGTGATCAAAAGCGTTATTCTGCAGCAGCAGTGTAAACATAATGGT
GAATATCTGACCGCCAGCGAAGTGCATAATTACATTAAACGCTATATGTATCAGTGCCGCAGCAATATGAAACCGCTGAGCTGTAAAGT
TATTGTGGCAGGTATTAATCCGGATGGCAGCAAATTTCTGGCATGTACCGATCCGTATGGTGCAAGCTGGGAAAGCGATCACATTGGCA
CCGGTTTTGGTAAATATCTGCAGGGTCTGCAGATTGCCGATGTTGTTAATGGTAGCTTTGATGATGTGAAAAAGGGCATCACCGAAGTT
TTTCGTGCAGTTAATGCCCGTAATACCACCGCAAATGGTAAAATCGAATTTATCACCGTTACACCGCAGGGCATTAATCATCTGGCACC
GGAACAAATTGATCCGAATTGGGAAGTTGTTGAAGGCACCTGGGATCAGAGCGCTTGGAGCCACCCGCAGTTCGAAAAAGGTGGAGGTT
CTGGCGGTGGATCGGGAGGTTCAGCGTGGAGCCACCCGCAGTTCGAGAAATAAgcggccgc

Insert BamHI_Ump-1_NotI

ggatccGCCACCATGTACGAACAGTGGATTCCGGAAGAACCGGTTGATCGTCTGCGTGAAGGTCTGCCGAATCTGCGTCATGGTAGCGT
TAATCAGCATCCGCTGGAAATTGCCATTGAAGAACGTCGTAAAACCCAGTTCAAAGACAAATTTGATGAACTGGCACTGCTGTATGGTG
AAGGTTTTGCAAATCATGAGAAGATGCTGTACAAACTGATTAAAAGCACCCGTATTGGTTTTCGCACCTATGAACGTCCGGATGATCTG
GCAATTGAAGTTTTTACCGGTGATATCGATGATGTGGATTTCAACGATATGTTTGCACCGAATGGTATGCGTGCCGATATTGAATTTGA
TCCGCATGAAATCCAAGAGAAACGCCTGAATATTGAATAAgcggccgc

Fig. S2. **Cryo-EM workflow of data processing** The image processing workflow employed for reconstructing the Tv20S structures in Cryo-Electron Microscopy (Cryo-EM). Representative images of data from **a** Tv20S-MZB dataset and from **b** Tv20S-CP-17. Both datasets were acquired at Titan Krios with Falcon 4i detector under identical conditions using the same setup (refer to Table S1 for details). In scheme **c** is an outline of Cryo-EM workflow and illustrative details from processing of each dataset. The ab-initio model served as the starting point for homologous refinement (Homo refine) to enhance the quality of the maps. Multiple iterations, including 3D classification and 2D classification, were carried out in several rounds to eliminate unwanted particles, refine the resolution, and improve the maps. The unsharpened maps of the final reconstruction and the gold-standard FSC curve using different masks are shown. The figures of the maps were generated by ChimeraX.[1]

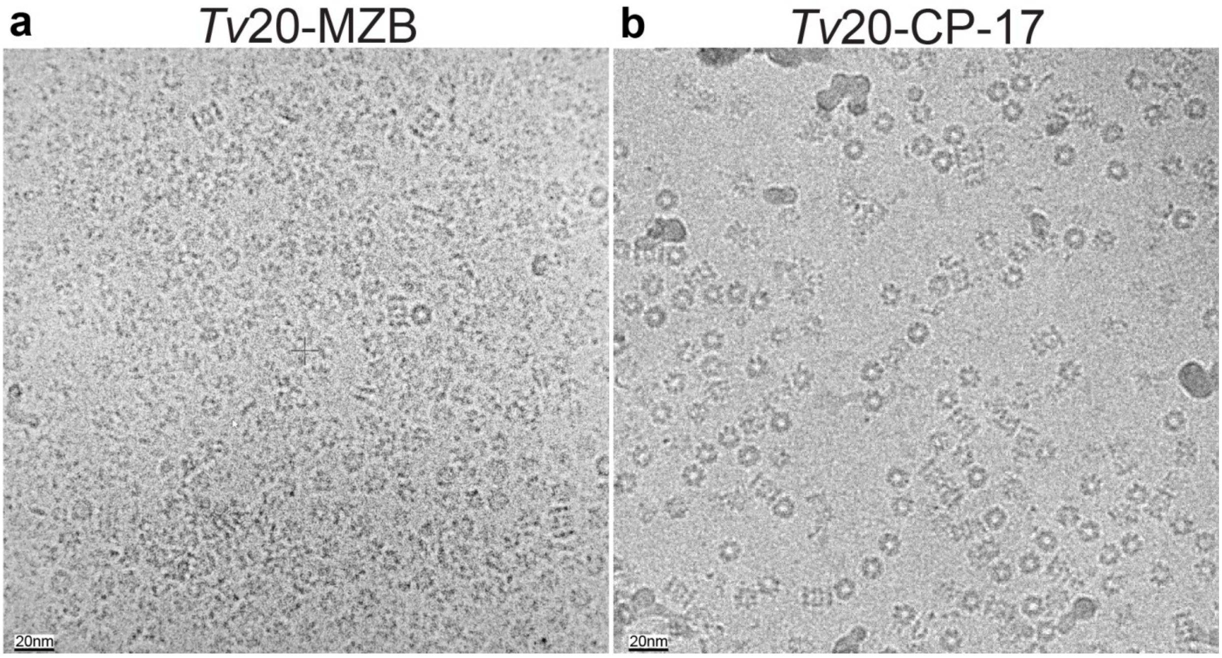

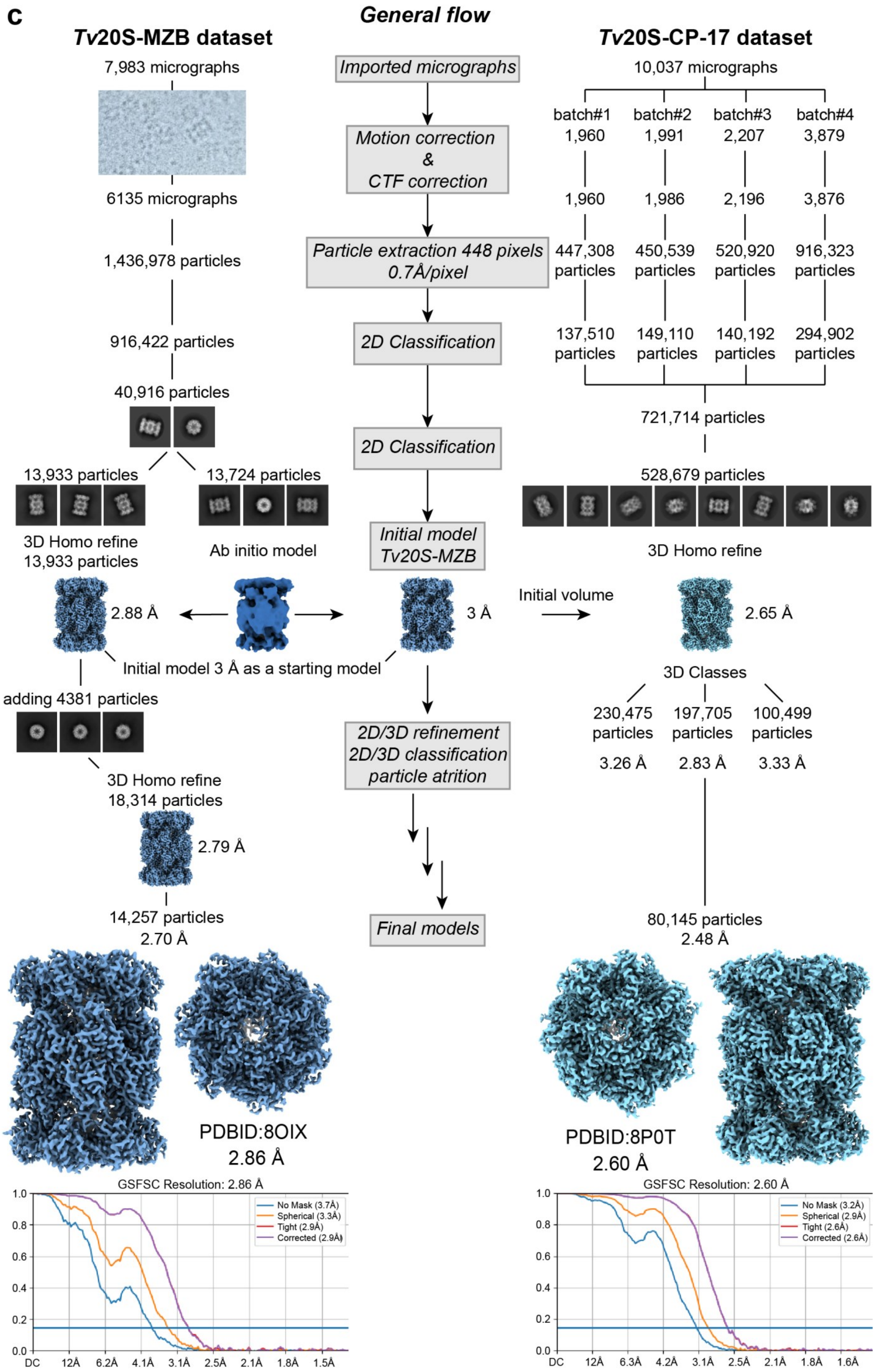
c
General flow
Tv20S-MZB dataset
Tv20S-CP-17 dataset
Imported micrographs
Motion correction & CTF correction
Particle extraction 448 pixels 0.7Å/pixel
2D Classification
2D Classification
Initial model Tv20S-MZB
2D/3D refinement 2D/3D classification particle atrition
Final models
7,983 micrographs
6135 micrographs
1,436,978 particles
916,422 particles
40,916 particles
13,933 particles
13,724 particles
3D Homo refine 13,933 particles
Ab initio model
2.88 Å
3 Å
Initial model 3 Å as a starting model
adding 4381 particles
3D Homo refine 18,314 particles
2.79 Å
14,257 particles 2.70 Å
PDBID:8OIX 2.86 Å
10,037 micrographs
batch#1 1,960
batch#2 1,991
batch#3 2,207
batch#4 3,879
1,960
1,986
2,196
3,876
447,308 particles
450,539 particles
520,920 particles
916,323 particles
137,510 particles
149,110 particles
140,192 particles
294,902 particles
721,714 particles
528,679 particles
3D Homo refine
Initial volume
2.65 Å
3D Classes
230,475 particles 3.26 Å
197,705 particles 2.83 Å
100,499 particles 3.33 Å
80,145 particles 2.48 Å
PDBID:8P0T 2.60 Å
GSFSC Resolution: 2.86 Å
No Mask (3.7Å)
Spherical (3.3Å)
Tight (2.9Å)
Corrected (2.9Å)
GSFSC Resolution: 2.60 Å
No Mask (3.2Å)
Spherical (2.9Å)
Tight (2.6Å)
Corrected (2.6Å)

**Table S1. Cryo-EM statistics for data collection, refinement and validation statistics**

| Deposited Cryo-EM structure | **Tv20S-MZB**<br>PDBID:8OIX | **Tv20S-CP-17**<br>PDBID:8P0T |
|---|---|---|
| Microscope | Titan Krios | Titan Krios |
| Detector | Falcon IVi | Falcon IVi |
| Magnification (nominal) | 165,000x | 165,000x |
| Voltage (kV) | 300 | 300 |
| Spherical aberration | 2.7 mm | 2.7 mm |
| Total electron dose ($e^-$) | 40 | 40 |
| Defocus range (µm) | (-2.4)–(-0.8) | (-2.4)–(-0.8) |
| Exposure (s) | 2.25 | 2.25 |
| Pixel size (Å) | 0.7 | 0.7 |
| Number of Micrographs | 7,983 | 10,037 |
| Final number of particles | 14,257 | 80,145 |
| Map resolution (Å)<br>[FSC threshold] | 2.86 [$FSC_{0.143}$] | 2.60 [$FSC_{0.143}$] |

**Refinement**

| | | |
|---|---|---|
| Initial model used (PDBID) | 7ZYJ | 8OIX |
| Validation | | |
| MolProbity score | 2.22 | 2.39 |
| Clashscore, all-atom | 24.83 | 28.72 |
| Rotamer outliers | 2.41% | 3.84% |
| Ramachandran plot | | |
| Favoured | 97.83% | 98.19% |
| Outliers | 0.00% | 0.05% |

**Table S2. Structural Alignment of 20S Proteasome Tv20-CP1 (PDB: 7ZYJ) with Leishmania and Human 20S Proteasomes, Including Percentage Identity of All Subunits between Human and *T. vaginalis***

The table of RMSD values for alignments of Cα of indiviudal chains Tv20-CP17 with Leishmania tarentolae 20S proteasome and human proteasome:

| | RMSD Leismania 7ZYJ | Chain ID | RMSD **Human 7PG9** | % Identity Human vs *T. vaginalis* | | RMSD Leismania 7ZYJ | Chain ID | RMSD **Human 7PG9** | % Identity Human vs *T. vaginalis* |
|---|---|---|---|---|---|---|---|---|---|
| **α1** | 1.252 | A | **1.153** | 42.08 | **β1** | 0.975 | H | **0.976** | 40.19 |
| **α2** | 1.219 | B | **1.031** | 50.22 | **β2** | 0.807 | I | **0.989** | 44.09 |
| **α3** | 0.985 | C | **1.260** | 49.20 | **β3** | 0.886 | J | **0.898** | 51.22 |
| **α4** | 1.048 | D | **1.028** | 49.79 | **β4** | 0.960 | K | **1.289** | 40.84 |
| **α5** | 1.252 | E | **1.099** | 51.04 | **β5** | 0.870 | L | **1.010** | 42.69 |
| **α6** | 1.116 | F | **1.144** | 45.06 | **β6** | 1.228 | M | **1.050** | 35.27 |
| **α7** | 1.121 | G | **1.205** | 41.67 | **β7** | 1.178 | N | **1.489** | 25.82 |

Overall RMSD values:
20S proteasome Tv20-CP17 with Leishmania tarentolae 20S proteasome **RMSD = 2.729**
20S proteasome Tv20-CP17 with human 20S proteasome **RMSD = 2.541**
The average RMSD values:
for all chains of Lesmania tarentolae 20S proteasome RMSD = 1.064
20S proteasome Tv20-CP17 with human 20S proteasome RMSD = 1.116*
(RMSD values were calculated for Tv20-CP17 (8P0T as a fix model) and human crystal structure (7PG9) for Cα atoms in the Pymol [The PyMOL Molecular Graphics System, Version 2.5.5 Schrödinger, LLC] using the command:align)

Fig. S3**. Sequence alignment of Tv20S and human 20S proteasomes and Ump-1 chaperone**

Sequence Alignment of α, β and Ump-1 subunits from *Trichomonas vaginalis* 20S proteasome and comparison with human (Hs) subunits. a) Multiple sequence alignment of all seven α subunits (α1-α7) from Tv20S. with their corresponding human α subunits (Hs20S). Conserved residues are boxed, and identical residues are highlighted in red. **b)** Multiple sequence alignment of all seven β subunits (β1-β7) Tv20S. Conserved residues are boxed, and identical residues are highlighted in red. **c)** Individual sequence alignment of each α subunit from Tv20S with their corresponding human α subunits (Hs20S). Conservation patterns and variations between species are illustrated. **d)** Individual sequence alignment of each β subunit from Tv20S with their corresponding human α subunits (Hs20S). Conservation patterns and variations between species are illustrated **e)** Single sequence alignment of Tv and Hs chaperone Ump-1, highlighting conserved regions. **f)** Sequence alignment of beta 2 active subunit from *Trichomonas vaginalis (Tv), Giardia lamblia (Gl), Plasmodium falciparum (Pf), Trypanosoma brucei (Tb), Homo sapiens (Hs)*, and *Ixodes ricinus (Ir)*. The asterisk denotes highly conserved residues Lys33 and a loop consisting of Ala46, Ala49, and Asn52. The alignment was done according to Robert, X. and Gouet, P. (2014).[2]

a

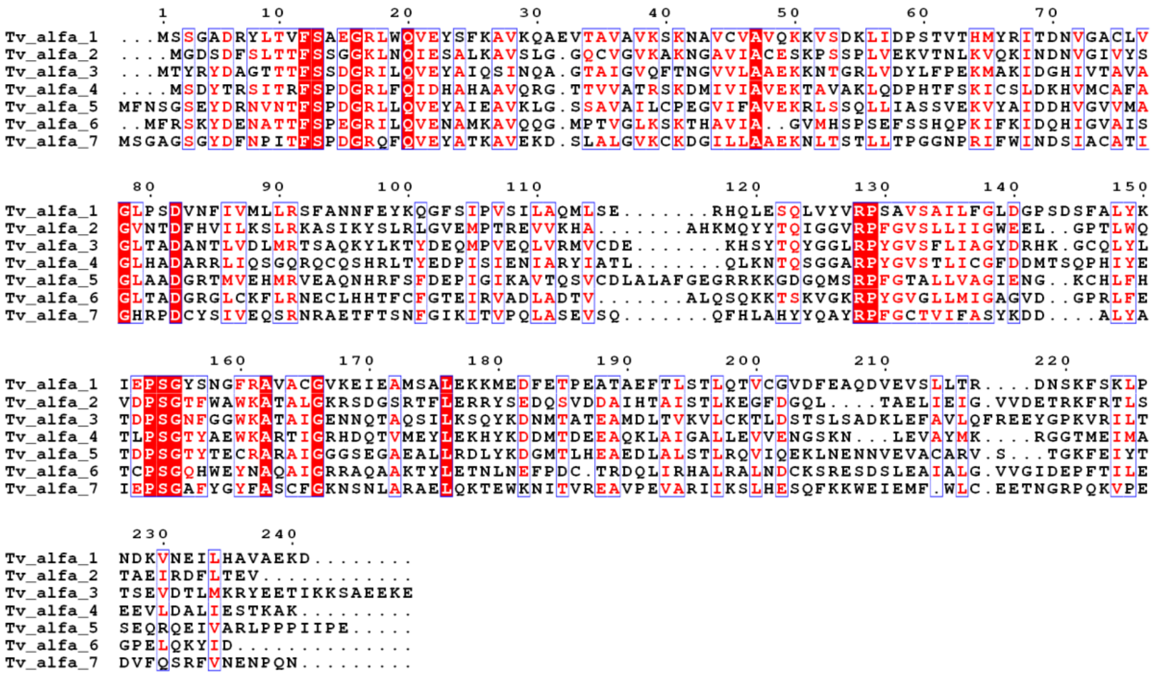

b

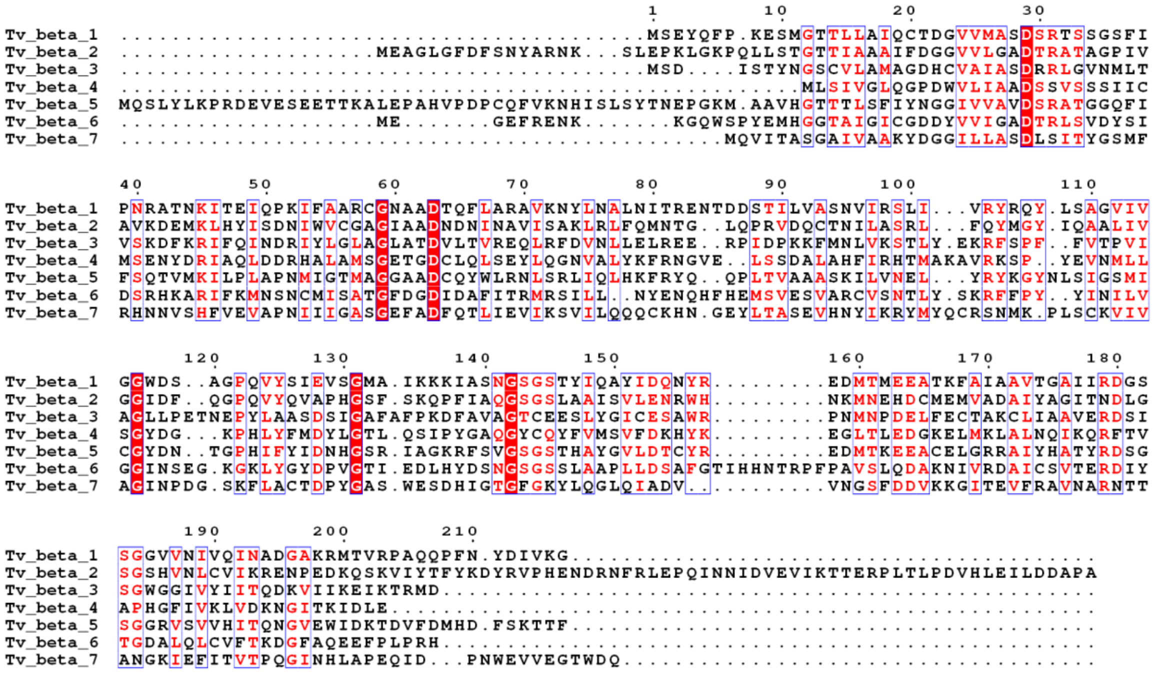

c

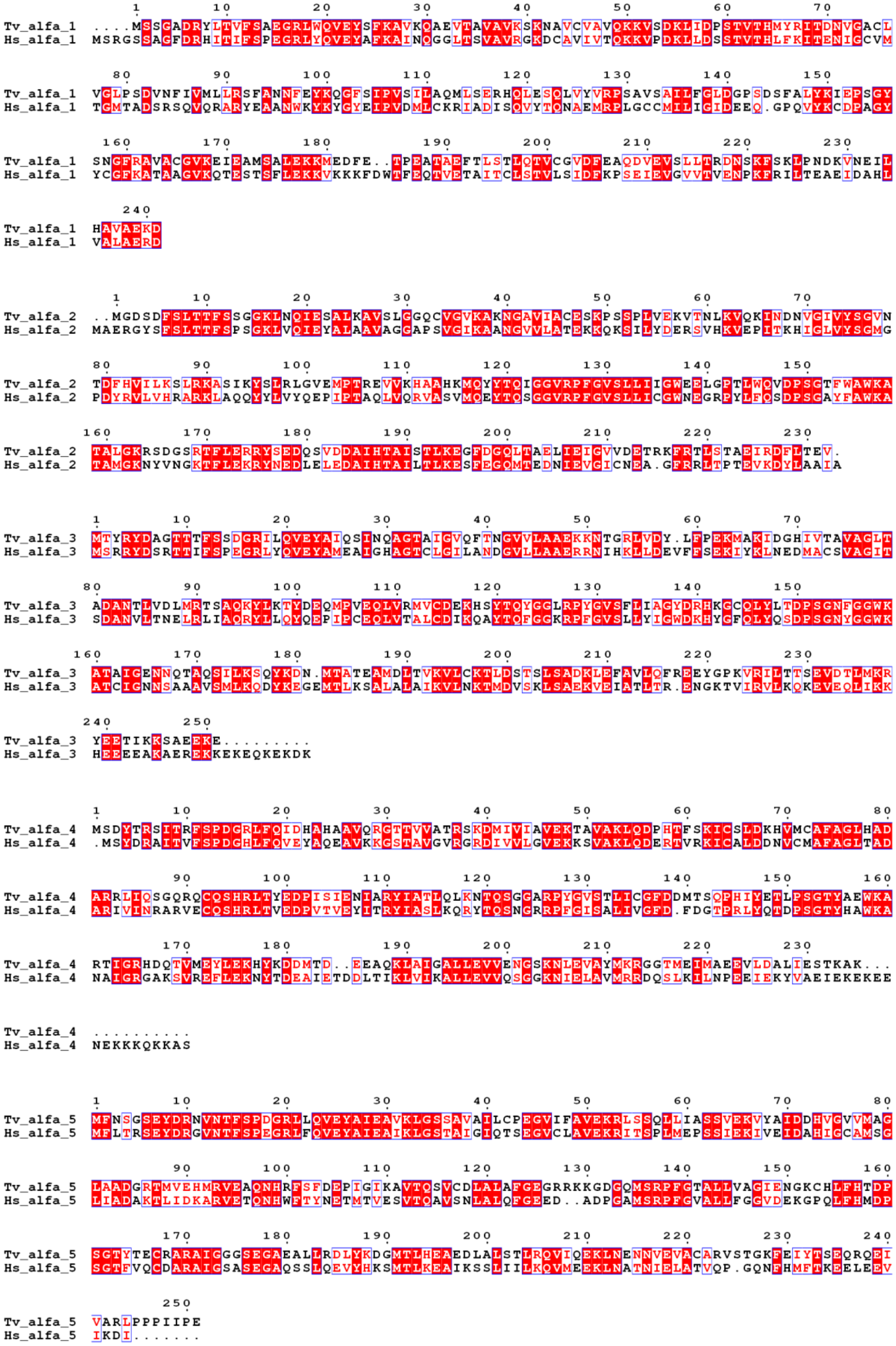
Tv_alfa_1
Hs_alfa_1
Tv_alfa_2
Hs_alfa_2
Tv_alfa_3
Hs_alfa_3
Tv_alfa_4
Hs_alfa_4
Tv_alfa_5
Hs_alfa_5

```
Tv_alfa_6  MFRSKYDENATTFSPEGRILQVENAMKAVQQGMPTVGLKSKTHAVIAGVMHSPSEFSSHQPKIFKIDQHIGVAISGLTAD
Hs_alfa_6  MFRNQYDNDVTVWSPQGRIHQIEYAMEAVKQGSATVGLKSKTHAVLVALKRAQSELAAHQKKILHVDNHIGISIAGLTAD

Tv_alfa_6  GRGLCKFLRNECLHHTFCFGTEIRVADLADTVALQSQKKTSKVGKRPYGVGLLMIGAGVDGPRLFETCPSGQHWEYNAQA
Hs_alfa_6  ARLLCNFMRQECLDSRFVFDRPLPVSRLVSLIGSKTQIPTQRYGRRPYGVGLLIAGYDDMGPHIFQTCPSANYFDCRAMS

Tv_alfa_6  IGRRAQAAKTYLETNLNEFPDCTRDQLIRHALRALNDCKSRESD.SLEAIALGVVGIDEPFTILEGPELQKYID......
Hs_alfa_6  IGARSQSARTYLERHMSEFMECNLNELVKHGLRALRETLPAEQDLTTKNVSIGIVGKDLEFTIYDDDDVSPFLEGLEERP

Tv_alfa_6  ........................
Hs_alfa_6  QRKAQPAQPADEPAEKADEPMEH


Tv_alfa_7  MSGAGSGYDFNPITFSPDGRQFQVEYATKAVEKDSLALGVKCKDGILLAAEKNLTSTLLTPGGNPRIFWINDSIACATIG
Hs_alfa_7  MSSIGTGYDLSASTFSPDGRVFQVEYAMKAVENSSTAIGIRCKDGVVFGVEKLVLSKLYEEGSNKRLFNVDRHVGMAVAG

Tv_alfa_7  HRPDCYSIVEQSRNRAETFTSNFGIKITVPQLASEVSQQFHLAHYYQAYRPFGCTVIFASYK...DDALYAIEPSGAFYG
Hs_alfa_7  LLADARSLADIAREEASNFRSNFGYNIPLKHLADRVAMYVHAYTLYSAVRPFGCSFMLGSYSVNDGAQLYMIDPSGVSYG

Tv_alfa_7  YFASCFGKNSNLARAELQKTEWKNITVREAVPEVARIIKSLHES.QFKKWEIEMFWLCEETNGRPQKVPEDVFQSRFV..
Hs_alfa_7  YWGCAIGKARQAAKTEIEKLQMKEMTCRDIVKEVAKIIYIVHDEVKDKAFELELSWVGELTNGRHEIVPKDIREEAEKYA

Tv_alfa_7  NENPQN.........
Hs_alfa_7  KESLKEEDESDDDNM
```

**d**

```
Tv_beta_1  ......................MSEYQFPKESMGTTLLAIQCTDGVVMASDSRTSSGSFIPNRATNKITEIQPKIFAARC
Hs_beta_1  MAATLLAARGAGPAPAWGPEAFTPDWESREVSTGTTIMAVQFDGGVVLGADSRTTTGSYIANRVTDKLTPIHDRIFCCRS

Tv_beta_1  GNAADTQFLARAVKNYLNALNITRENTDDSTILVASNVIRSLIVRYRQYLSAGVIVGGWDS.AGPQVYSIEVSGMAIKKK
Hs_beta_1  GSAADTQAVADAVTYQLGFHSIEL..NEPPLVHTAASLFKEMCYRYREDLMAGIIIAGWDPQEGGQVYSVPMGGMMVRQS

Tv_beta_1  IASNGSGSTYIQAYIDQNYREDMTMEEATKFAIAAVTGAIIRDGSSGGVVNIVQINADGAKRMTVRPAQQPF.NYDIVKG
Hs_beta_1  FAIGGSGSSYIYGYVDATYREGMTKEECLQFTANALALAMERDGSSGGVIRLAAIAESGVERQVLLGDQIPKFAVATLPP

Tv_beta_1  .
Hs_beta_1  A


Tv_beta_2  .......MEAGLGFDFSNYARNKSLEPKL.....GKPQLLSTGTTIAAAIFDGGVVLGADTRATAGPIVAVKDEMKLHYI
Hs_beta_2  MAAVSVYAPPVGGFSFDNCRRNAVLEADFAKRGYKLPKVRKTGTTIAGVVYKDGIVLGADTRATEGMVVADKNCSKIHFI

Tv_beta_2  SDNIWVCGAGIAADNDNINAVISAKLRLFQMNTGLQPRVDQCTNILASRLFQYMGYIQAALIVGGIDFQGPQVYQVAPHG
Hs_beta_2  SPNIYCCGAGTAADTDMTTQLISSNLELHSLSTGRLPRVVTANRMLKQMLFRYQGYIGAALVLGGVDVTGPHLYSIYPHG

Tv_beta_2  SFSKQPFIAQGSGSLAAISVLENRWHNKMNEHDCMEMVADAIYAGITNDLGSGSHVNLCVIKRENPEDKQSKVIYTFYKD
Hs_beta_2  STDKLPYVTMGSGSLAAMAVFEDKFRPDMEEEEAKNLVSEAIAAGIFNDLGSGSNIDLCVISKNKL.........DFLRP

Tv_beta_2  YRVPHENDRNF...R.........LEPQINNIDVEVIKTTERPLTLPDVHLEILDDAPA
Hs_beta_2  YTVPNKKGTRLGRYRCEKGTTAVLTEKITPLEIEVLEETVQTMDTS............


Tv_beta_3  MSDISTYNGSCVLAMAGDHCVAIASDRRLGVNMLTVSKDFKRIFQINDRIYLGLAGLATDVLTVREQLRFDVNLLELREE
Hs_beta_3  .MSIMSYNGGAVMAMKGKNCVAIAADRRFGIQAQMVTTDFQKIFPMGDRLYIGLAGLATDVQTVAQRLKFRLNLYELKEG

Tv_beta_3  RPIDPKKFMNLVKSTLYEKRFSPFFVTPVIAGLLPETNEPYLAASDSIGAFAFPKDFAVAGTCEESLYGICESAWRPNMN
Hs_beta_3  RQIKPYTLMSMVANLLYEKRFGPYYTEPVIAGLDPKTFKPFICSLDLIGCPMVTDDFVVSGTCAEQMYGMCESLWEPNMD

Tv_beta_3  PDELFECTAKCLIAAVERDSISGWGGIVYIITQDKVIIKEIKTRMD
Hs_beta_3  PDHLFETISQAMLNAVDRDAVSGMGVIVHIIEKDKITTRTLKARMD
```

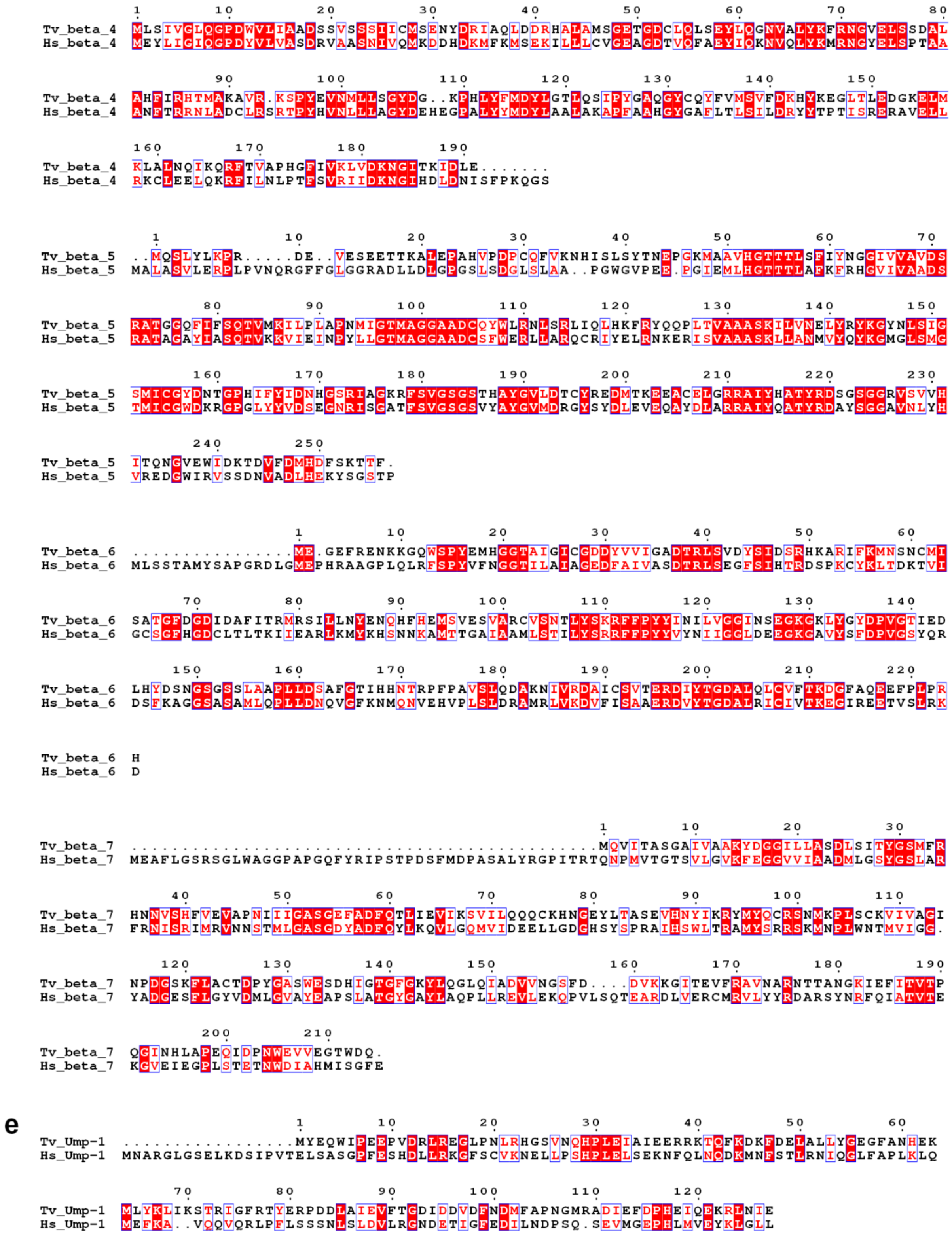

**f**

```
           1        10        20        30        40        50        60        70        80
           .        .         .         .         .         .         .         .         .
Tv_beta_2  TTIAAAIFDGGVVLGADTRATAGPIVAVKDEMKLHYISDNIWVCGAGIAADNDNINAVISAKLRLFQMNTGLQPRVDQCT
Gl_beta_2  TTIMGLTFKGGVILAADTRSTGGPVVMNKNKKKLVCINEQMYMAGAGTAADTAAVGRMAASSLRLHAYKTGRKPLVQSAV
Pf_beta_2  TTICGLVCQNAVILGADTRATEGPIVADKNCSKLHYISKNIWCAGAGVAGDLEHTTLWLQHNVELHRLNTNTQPRVSMCV
Tb_beta_2  TTIVGVVFEGGVVLGADTRATEGSIVADKRCKKIHYMAPNIMCCGAGTAADTEAVTNMTAANLTLHRLGTGKQSRVSEAL
Hs_beta_2  TTIAGVVYKDGIVLGADTRATEGMVVADKNCSKIHFISPNIYCCGAGTAADTDMTTQLISSNLELHSLSTGRLPRVVTAN
Ir_beta_2  TTIAGIIFKDGVILGADTRATSGSIIADKNCAKIHYMAPNIYCCGAGTAADTEKTTNMISSQLELHRLATGRTVPVCTAN
                                            *            *  * *

                   90        100       110       120       130       140       150       160
                   .         .         .         .         .         .         .         .
Tv_beta_2  NILASRLFQYMGYIQAALIVGGIDFQGPQVYQVAPHGSFSKQPFIAQGSGSLAAISVLENRWHNKMNEHDCMEMVADAIY
Gl_beta_2  KLISDHLFRYMGYVSAYVIIAGVDYKGAHLCSIDAHGSITAQPFVTDGSGRLAAQSMLEKHWKPDMEENDAVECACKAVE
Pf_beta_2  SRLTQELFKYQGYKVCAIVLGGVDVNGPQLYGIHPHGSSCLLPFTALGSGSLNAMAVLEAKYRDNMTIEEGKNLVCEAIC
Tb_beta_2  TLLKRHLYRYQGHVSAALVLGGVDVGGAFLATVAPHGSTDRLPFVAMGSGSIAAMAALETGYKENLTLEEAKQLVVSAIH
Hs_beta_2  RMLKQMLFRYQGYIGAALVLGGVDVTGPHLYSIYPHGSTDKLPYVTMGSGSLAAMAVFEDKFRPDMEEEEAKNLVSEAIA
Ir_beta_2  RMLKQMLFRYQGHISAALVLGGVDNTGGVLCCIHPHGSTDRLPYVTMGSGSLAAMAVFERDWKPDMTLEEGKKVVRDAIA

                   170       180       190       200            210       220       230
                   .         .         .         .              .         .         .
Tv_beta_2  AGITNDLGSGSHVNLCVIKRENPEDKQSKVIYTFYKDYRVPHENDRN....FRLEPQINNIDVEVIKTTERPLTLPDVHL
Gl_beta_2  AGVFNDLGSGSGVDLVILRLDGTSELKHCIRESGSYSGSAPTGVYY..................................
Pf_beta_2  AGIFNDLGSGGNVDICVITKDSYQHIRPYKEPNMRL.YHLPHPTIYPKGTTPILSEKIEYIKKFISVEDA..........
Tb_beta_2  KGIFNDPYSGTQVDLCVITKAKTEMLIGYDKPNDRK.YPK.HDIKLPPGTTPILREEIRQLVTITELE............
Hs_beta_2  AGIFNDLGSGSNIDLCVISKNKLDFLRPYTVPNKKG.TRL.GRYRCEKGTTAVLTEKITPLEIEVLEETVQTMD.....T
Ir_beta_2  AGIFNDLGSGSNIDICVITREGKEYIRPYDEANKKG.QRQ.LSYKFKKGTTAVLSSNVVPIEVEET..TVRHLD.....A

            240
            .
Tv_beta_2  EILDDAPA
Gl_beta_2  ........
Pf_beta_2  ........
Tb_beta_2  ........
Hs_beta_2  .....S..
Ir_beta_2  EAMDTSA.
```

Fig. S4. **Structural comparison of Cryo-EM maps of the inhibitor molecules MZB and CP-17 covalently bound in the active sites of *Tv*20S**

The top views of both Tv20S structures with **a** MZB and **b** CP-17 inhibitors (both shown as blue sticks). Cryo-EM Maps are carved around inhibitors for clarity as green mesh at sigma 4. Detailed views on the active sites are shown as cartoon and sticks for inhibitors. MZB (panels c,d and e) and CP-17 (panels f, g and) are displayed with active sites in order: β1 (panels c&f are shown in yellow), β2 (panels d&g, in cyan), and β (panel e&h, magenta).

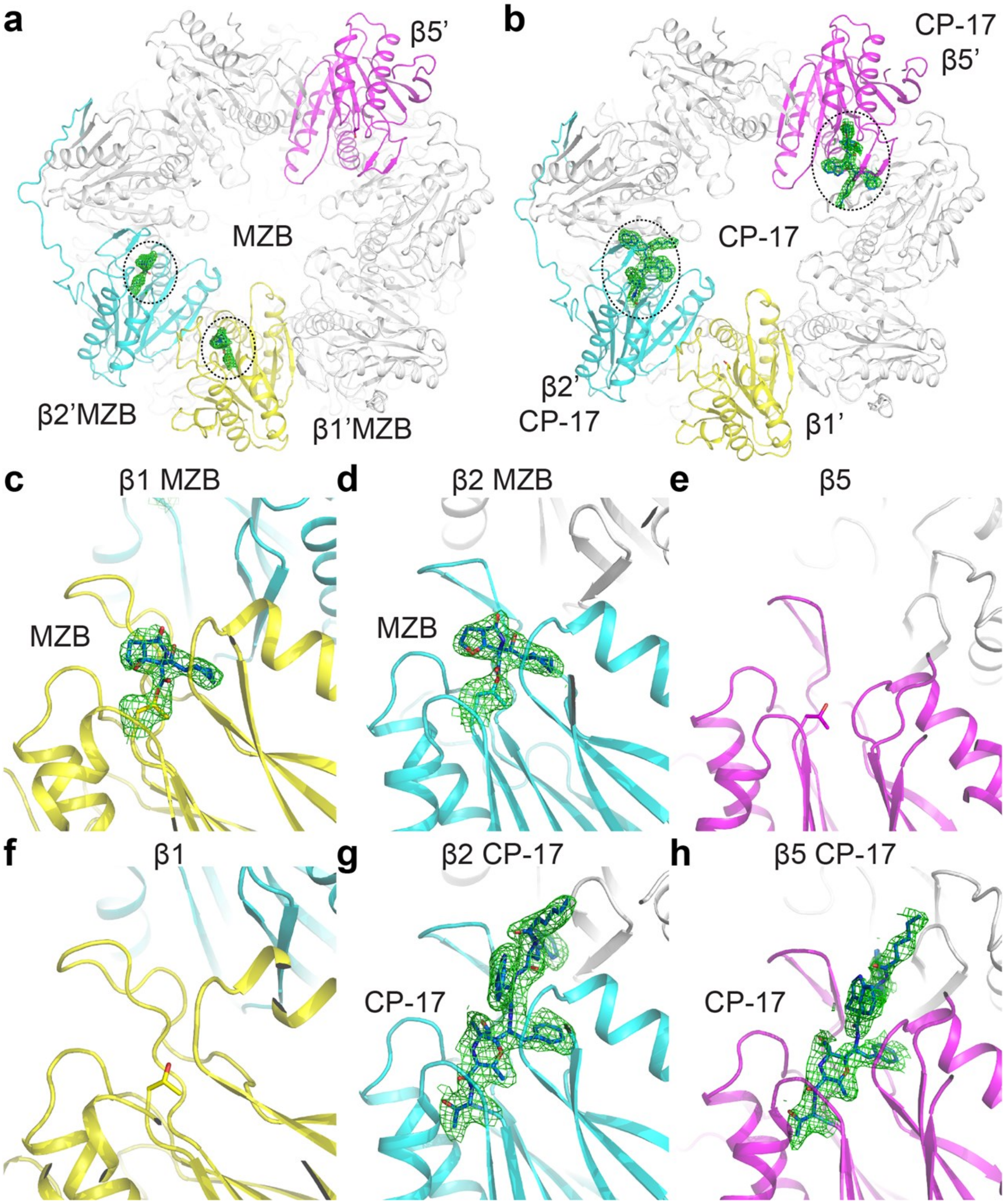

Fig. S5. **Cryo-EM maps in all three active sites for both *Tv*20S structures**

Comparison of cryo-EM maps for both Tv20S structures with MZB (**a**,**b**,**c**) and CP-17 (**d**,**e**,**f**) inhibitors. Maps are shown as gray mesh at sigma 4 and density around inhibitors is depicted in green. Active sites are displayed in order: β1 (panels a&d are shown in yellow), β2 (panels **b**&**e**, in cyan), and β (panel **c**&**f**, in magenta).

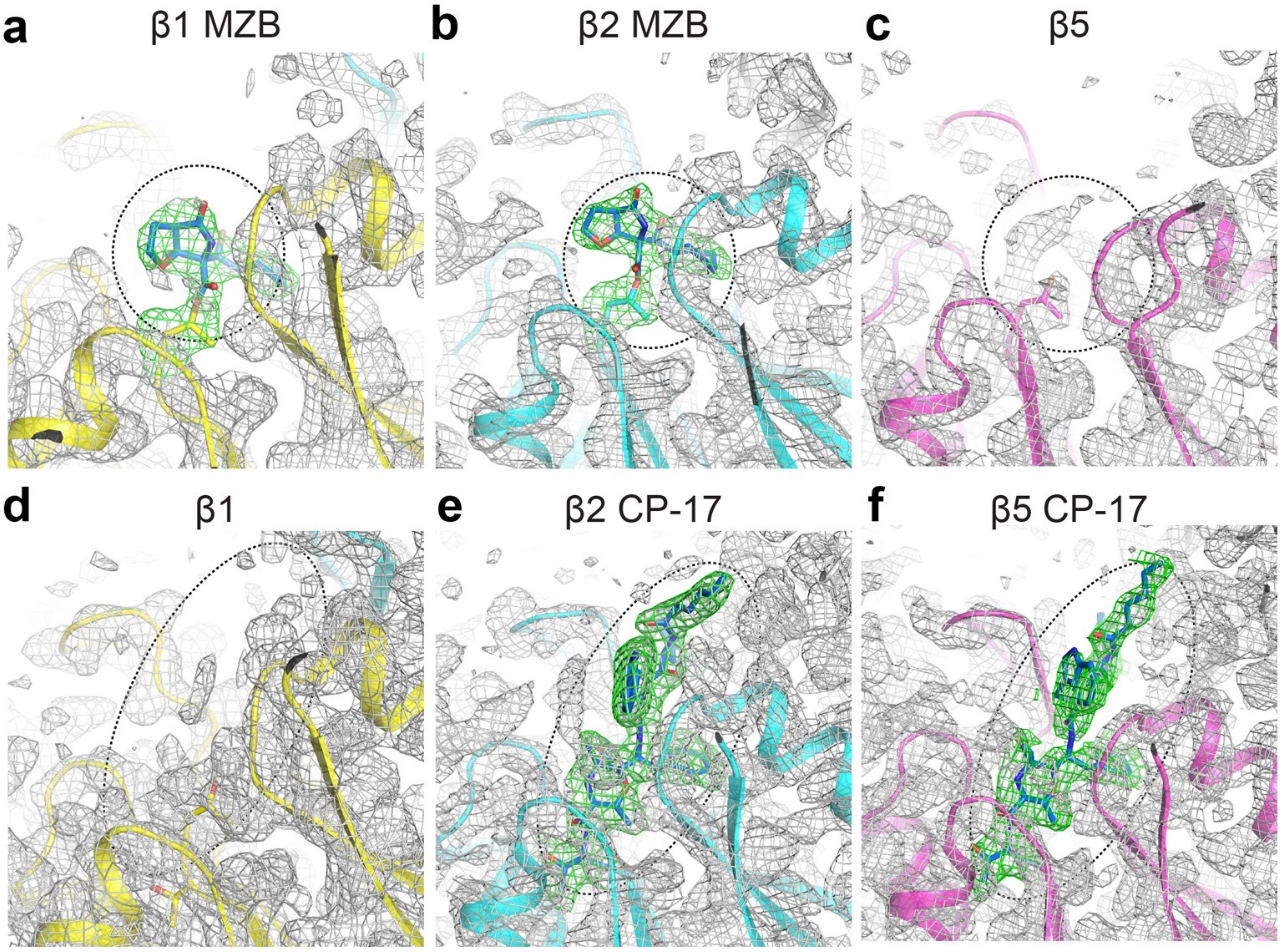

Fig. S6. **Prospects for developing inhibitors that target Cys46 of the β1 subunit of *Tv*20S proteasome**

**a** The active site of the β1 subunit in the Tv20S-MZB proteasome structure is highlighted in yellow, while MZB is shown as blue sticks. This region includes neighbouring residues, including Cys46. **b** The structural overlay of β1 and β2 (cyan sticks) subunits of Tv20S-MZB proteasome. **c** An overlay comparison is made between the β1 active site of Tv20S-MZB (shown in yellow, with MZB represented as blue sticks) and the human 20S structure, PDB ID = 7PG9 (shown in orange). **d** Another overlay is presented, showcasing only the CP-17 inhibitor (depicted in white) of the β2 subunit onto the β1 active site of Tv20S-MZB (represented in yellow, with MZB shown as blue lines).

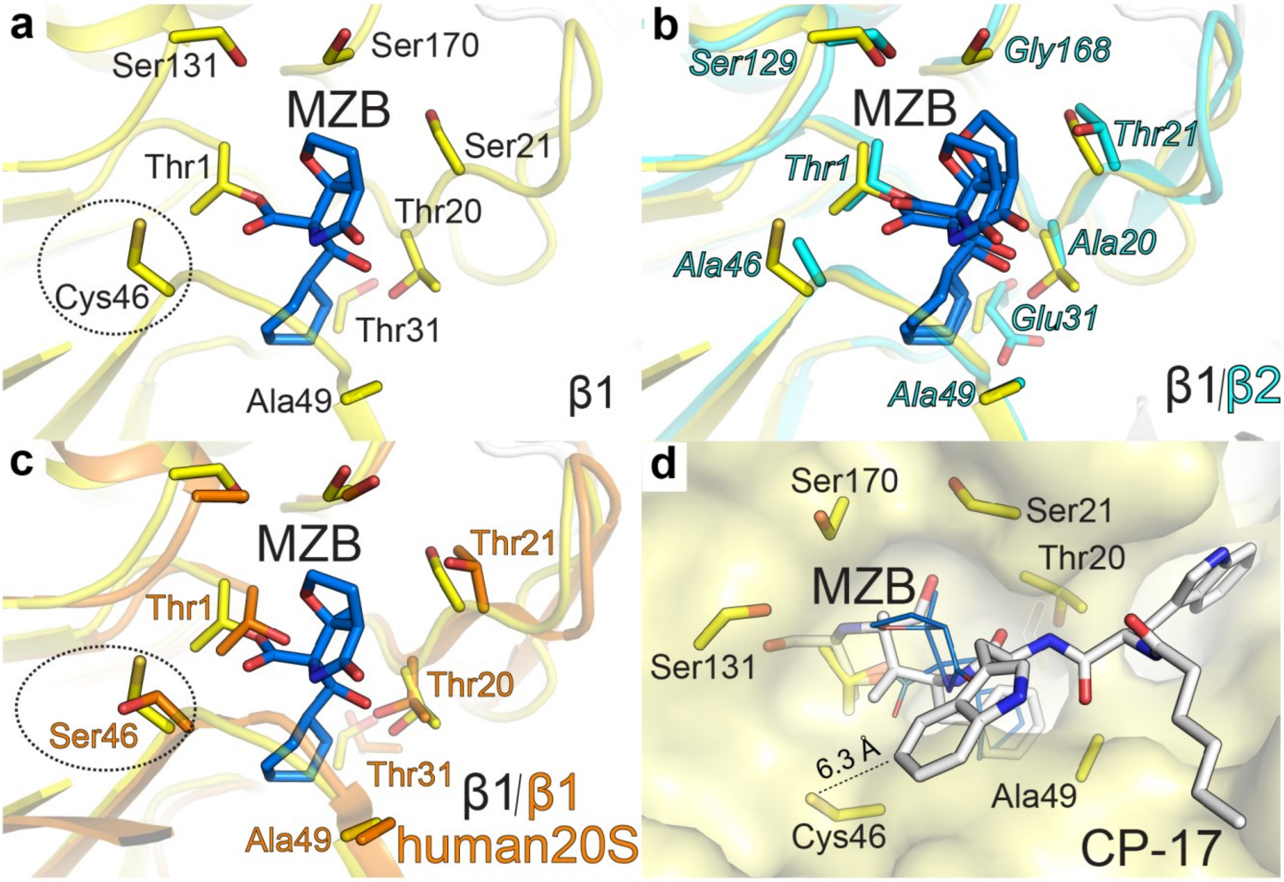

Fig. S7. **Model of CP-17 "virtual" clashes within the *Tv*20S proteasome β1 active site pocket**

**a** The β1 active site of Tv20S-MZB structure (shown in yellow, MZB in blue) with neighbouring residues including Cys46. **b** The surface of β1 with MZB (yellow and blue sticks) and β2 with and overlay of surface and CP-17 inhibitor (white and white sticks) from the β2 of Tv20S-CP-17 proteasome. CP-17 is shown as white sticks for clarity. **c** Detail of β2 with CP-17 with highlighted residues mainly surrounding indole rings. **d** The inset of the figure shows surface of β2 S3 pocket of Tv20S-CP-17 proteasome. The main panel shows surface of β1 site with CP-17 (white) modelled by superimposition of β2 and β1 subunits. The β1 pocket is lined with much bulkier residues such as Pro27 and polar residues Ser118, Gln112 and Gln127. Unlike in the β2 pocket lined with smaller and more hydrophobic residues Ala (22,27,124,126,132) and Val28.

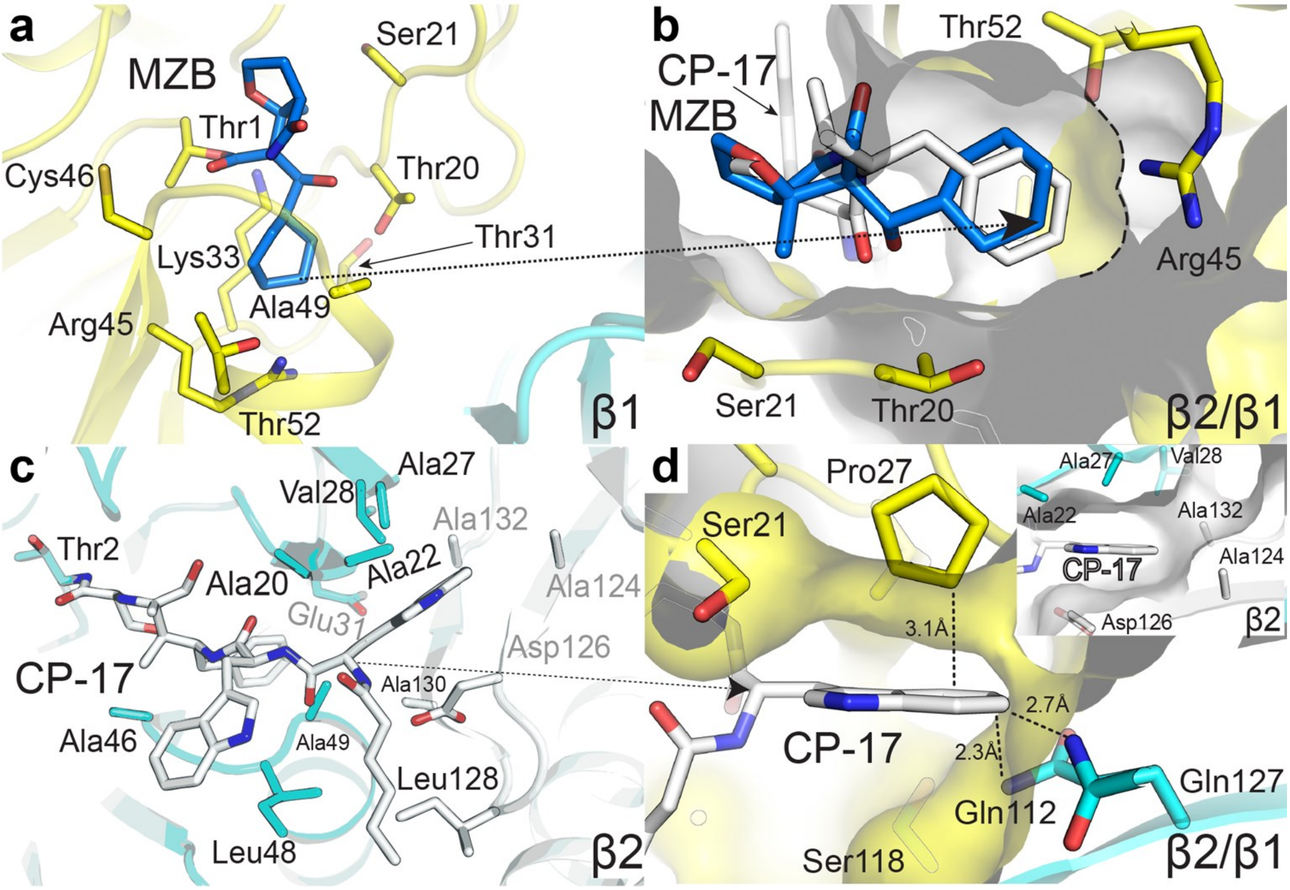

Fig. S8. **Inhibition of rTv20S by MZB and CP-17: $IC_{50}$ Values for β5, β2, and β1**

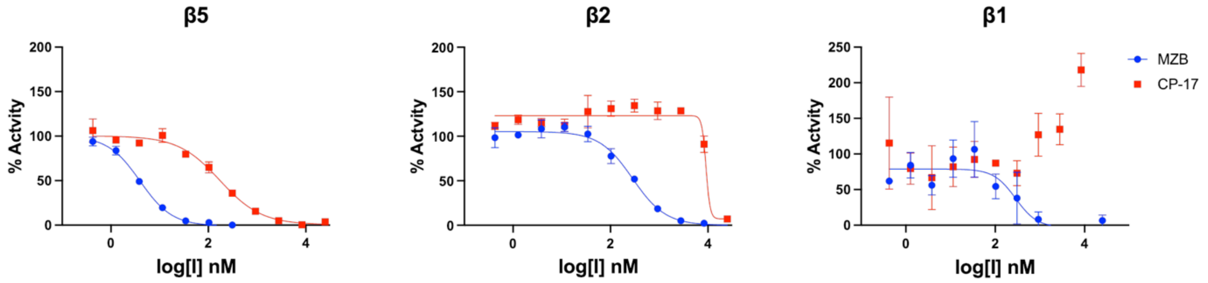


Inhibition of r*Tv*20S using MZB or CP-17 for β5, β2, and β1 subunits. The $IC_{50}$ values obtained after one hour of incubation of enzyme-inhibitor complex are represented as follows: for MZB: β5 = 3.94±0.24 nM, β2 = 283.70 ± 31.03nM and β1 = 284.40 ± 112.10 and for CP-17: β5 = 172.10 ± 22.36 nM, β2 = ~10 µM, β1 = no inhibition. The rate of AMC release was calculated in the presence of serial dilution of the inhibitor (25000.00 – 0.42 nM) and the $IC_{50}$ values were determined using GraphPad Prism 9 software.

**Suplementary references**

1 Pettersen, E. F. *et al.* UCSF ChimeraX: Structure visualization for researchers, educators, and developers. *Protein science : a publication of the Protein Society* **30**, 70-82 (2021). https://doi.org:10.1002/pro.3943

2 Robert, X. & Gouet, P. Deciphering key features in protein structures with the new ENDscript server. *Nucleic Acids Res* **42**, W320-324 (2014). https://doi.org:10.1093/nar/gku316
